# Oarfish: Enhanced probabilistic modeling leads to improved accuracy in long read transcriptome quantification

**DOI:** 10.1101/2024.02.28.582591

**Authors:** Zahra Zare Jousheghani, Rob Patro

**Affiliations:** Department of Electrical and Computer Engineering, University of Maryland, College Park, 20742, Maryland, USA; Department of Computer Science, University of Maryland, College Park, 20742, Maryland, USA

**Keywords:** RNA-seq, transcript quantification, long reads, Oxford Nanopore, PacBio

## Abstract

**Motivation:** Long read sequencing technology is becoming an increasingly indispensable tool in genomic and transcriptomic analysis. In transcriptomics in particular, long reads offer the possibility of sequencing full-length isoforms, which can vastly simplify the identification of novel transcripts and transcript quantification. However, despite this promise, the focus of much long read method development to date has been on transcript identification, with comparatively little attention paid to quantification. Yet, due to differences in the underlying protocols and technologies, lower throughput (i.e. fewer reads sequenced per sample compared to short read technologies), as well as technical artifacts, long read quantification remains a challenge, motivating the continued development and assessment of quantification methods tailored to this increasingly prevalent type of data.

**Results:** We introduce a new method and software tool for long read transcript quantification called `oarfish`. Our model incorporates a novel and innovative coverage score, which affects the conditional probability of fragment assignment in the underlying probabilistic model. We demonstrate that by accounting for this coverage information, `oarfish` is able to produce more accurate quantification estimates than existing long read quantification methods, particularly when one considers the primary isoforms present in a particular cell line or tissue type.

**Availability and Implementation:** `Oarfish` is implemented in the `Rust` programming language, and is made available as free and open-source software under the BSD 3-clause license. The source code is available at https://www.github.com/COMBINE-lab/oarfish.

## Introduction

Since the introduction of high-throughput RNA-sequencing [5, 28, 30, 31], the bioinformatics community has invested tremendous effort in the development of methods and software to tackle various challenges related to the analysis of this data. One of the first, and therefore one of the most fundamental challenges, is the accurate quantification of transcript and gene expression from this sequencing data — which has spurred the development of many methods for transcript quantification (e.g. [35, 24, 44, 16, 32, 38, 18, 7, 37] among many others).

The majority of these tools have been focused on increasing the accuracy or efficiency (or both) of quantification from high-throughput, short read data. Short reads have developed to have high base-level accuracy, small error rate, high reproducibility and immense throughput. Yet, they also have fundamental drawbacks that arise due to the relatively short read lengths used in sequencing (usually <350bp). This is considerably smaller than the average length of e.g. most spliced mRNA human transcripts (∼ 2Kbp [23]) that needed to be analyzed. A primary consequence of this mismatch between the length of sequenced reads and the length of the underlying transcripts that these reads are being used to measure is that the locus of origin of many of these reads remains fundamentally ambiguous. This problem is commonly called fragment ambiguity or read to transcript ambiguity [6], and it results in inferential uncertainty [39, 52] in the underlying transcript abundance estimates. That is, uncertainty in the transcript from which the sequenced reads originated leads to uncertainty in the estimated abundances of the transcripts being quantified, which, in turn, complicates and hampers downstream analyses. These uncertainties are somewhat reduced using common techniques such as paired-end sequencing [29, 6], but the problem remains, and is fundamental to such sequencing technologies.

While new computational methodologies have continued to advance and improve transcript quantification accuracy, they cannot overcome fundamental limitations inherent in the underlying measurements (i.e. sequenced data). However, the advent of the long read sequencing technologies, and their continued development in terms of reduced error rates and improved throughput, promises to mitigate or eliminate this fundamental limitation, and potentially revolutionized transcriptome analysis, as it has done with e.g. genome assembly [36]. Specifically, long reads have the ability to capture the whole length of the transcripts being sequenced within a single read, eliminating the potential ambiguity about the transcript from which the read has arisen. This reduction in (or elimination of) ambiguity has striking implications for both transcript discovery — that is, transcript assembly or identification — and transcript quantification. However, while the long reads can have increased specificity and overcome the ambiguity-related drawbacks of short read, they bring their own challenges, such as lower accuracy and higher error rate in comparison to short reads, and lower sequencing depth (even when sequencing a similar number of total nucleotides as a short-read sample, long reads comprise more nucleotides *per-read*, and so typically result in fewer independent measurements).

Currently, there are two major long read sequencing technologies; Oxford Nanopore Technologies (ONT) [1, 46, 48] and Pacific Biosciences (PacBio) [2, 40, 47], each technology with its own benefits and drawbacks. While the PacBio long reads have high accuracy and low error rate (especially HiFi reads), it typically has lower throughput and higher cost than ONT sequencing, though throughput can be improved substantially using techniques such as MAS-ISO-seq [3]. ONT sequencing typically provides higher throughput, and can therefore be more cost-efficient, but the generated reads typically have lower accuracy and higher error rate. Other differences in capabilities arise from the actual mechanisms by which sequencing is performed — for example, ONT sequencing is capable of sequencing either cDNA (as is traditionally used in RNA-seq) or performing direct RNA sequencing without first requiring reverse transcription, which also permits direct detection of base-level modifications of the underlying RNA molecule.

Long-read RNA sequencing is not without a growing collection of associated quantification methods. In response to the limitations of traditional tools, methods like TALON [49], FLAIR [42], and Mandalorion [9] have emerged, and while these tools support quantification, they primarily focus on novel transcript identification. For quantification, some tools initially developed for short-read data, like `salmon` [37] have since been augmented to support long-read data by adjusting the sequencing error models appropriately and removing the length dependence in the underlying graphical model, and recent benchmarks have shown that this approach works well for long read quantification [11]. At the same time, many tools have been developed specifically for use with long-read data, such as `Bambu` [11], `ESPRESSO` [15], `LIQA` [21], and `NanoCount` [17] among others. These tools also differ in the details of how they perform probabilstic read allocation. For example `Bambu`’s model applies an EM, but also categorizes reads into equivalence classes not just based on the transcripts to which they map, but on their *category* with respect to the target transcripts [11]. `ESPRESSO`’s latent variable model is dependent on the read-isoform compatibility matrix [15], and `LIQA`’s latent variable is influenced by the read quality score and the read length distribution [21] based on a survival model. The read allocation in `NanoCount` is connected to the number of alignments per read [17], and the method also applies several heuristic, but data driven, filters to remove alignemnts that are more likely than not do detract from quantification accuracy. Though these later long-read centric methods add interesting aspects to the quantification approach, they, too still tend to have a substantial focus on novel transcript discovery. While transcript discovery from long reads is an exciting and challenging problem, we argue here that even quantification with these data remain an unsolved challenge, and it deserves its own dedicated methods and approaches.

In this work, we focus primarily on quantification of ONT long reads, due to their higher throughput, lower cost, and the greater availability of publicly-available benchmarking datasets, all of which enable a broad range of usability of this technology. However, the models we develop and implement are likely largely technology agnostic, and therefore are likely to also improve the quantification accuracy for both major long read technologies. Likewise, reduced sequencing error and more accurate reads — if obtained at the same sequencing depth — are certainly key factors that we expect to contribute to more accurate quantification results and subsequently, more accurate and precise transcriptome analysis.

## Methods and Materials

In this section, we first review the popular quantification model [25, 44] used in previous short read quantification methods, and slightly adapted for long read quantification, then propose a novel addition to this model that can be used to improve quantification based on long read data (with the focus and evaluation here being on ONT data).

### Previous quantification framework

In [25], the authors propose a generative model for RNA-seq sequencing. A plate diagram of this generative model (which we do not reproduce here for lack of space) is provided in shown in Figure 1 of [25], and an extended model is presented in Figure 4 of [24]. The corresponding likelihood of a given collection of data (i.e. sequenced fragments and their corresponding alignments) is given as the following:

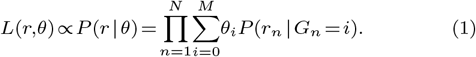

The likelihood defined in eq. (1) can be maximized (locally) using an Expectation-Maximization (EM) [13] algorithm, which the authors derive for this particular likelihood in [25, 24]. This algorithm is then used to obtain a maximum likelihood estimate (MLE) of the model’s parameters *θ* = [*θ*_0_, *θ*_1_,…, *θ*_*M*_], which correspond to the relative abundances of the isoforms, given the observed data *r* =[*r*_1_,*r*_2_,…,*r*_*N*_]. Here M is the number of isoforms and N is the number of fragments. In eq. (1), *M*_0_ represents a “noise” isoform, and *θ* _*i*_,*r*_*n*_,*G*_*n*_ parameters denote the relative abundance of *i*^th^ isoform, the *n*^th^ fragment, and the (latent) random variable assigning the *n*^th^ fragment to the *i*^th^ isoform. Therefore, *P* (*r*_*n*_ |*G*_*n*_ = *i*) is the conditional probability of observing the fragment *r*_*n*_ given that it arises from the *i*^*th*^ isoform.

**Fig. 1.**
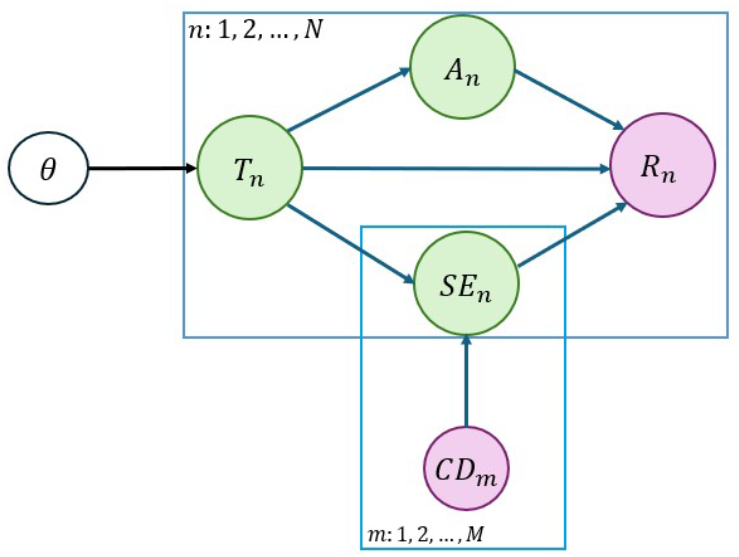
Graphical model for an improved generative model of RNA-seq long reads. θ represents the desired isoform expression level parameter. The latent variables *T*_*n*_, *A*_*n*_, and *SE*_*n*_ correspond to the isoform, alignment score, and start-end positions, respectively. The observed variables *R*_*n*_ and *CV*_*m*_ are associated with the read and coverage model, where *m* and *n* stand for the number of isoforms and reads, respectively. Green circles represent latent variables, white circles denote desired parameters, and pink circles indicate observed variables.

In [25, 24], an EM algorithm is applied to obtain a MLE estimate Θ, and this quantification model, or variants of it, have been used in many different methods designed for isoform expression quantification using short read RNA-seq data [24, 33, 34, 41] (and many others).

It is also worth highlighting that, shortly after Li et al. introduced RSEM [25], Turro et al. [44] introduced MMSeq. The latter tool is particularly notable because it introduced a related but distinct model in which likelihood was no longer evaluated (and parameters no longer optimized) at the level of individual fragments, but rather at the level of *equivalence classes* of fragments, where two fragments *f*_*i*_ and *f*_*j*_ are considered equivalent if and only if they map or align to the same set of transcript targets. By working with such equivalence classes, and retaining their multiplicity (i.e. the number of fragments belonging to each such class), MMSeq defined a similar but simplified likelihood which is *much* faster to optimize, as the E step of the EM algorithm scales with the number of distinct equivalence classes rather than with the number of distinct fragments. This alternative formulation of the problem represents an *approximate* factorization of the likelihood function, and has also since been used in several transcript quantification tools [44, 38, 51, 7, 37]. Between the extremes of the fragment-level model of Li et al. and the compatibility-based model of Turro et al. there exist a range of potential factorizations of the likelihood function, in which one can trade off computational efficiency for model fidelity [50], and factorizations can be derived that improve quantification accuracy relative to the compatibility-based model while still retaining rapid quantification speed.

In this work, we adopt a fragment-level likelihood, rather than a compatibility-based or alternative approximate likelihood factorization. Specifically, we take this approach as we will augment the model with an alignment-level term (a modification to the conditional fragment probability), which should be applied at the fragment, rather than equivalence class level. Moreover, given the typically smaller number of sequenced fragments in long read samples, the computational cost of adopting the fragment-level likelihood is relatively modest. Finally, we note that developing an appropriate approximate factorization that accounts for our modified model (or extensions of it) should be feasible, but as the current computational costs remain reasonable, we leave that to future work.

### Existing Long Read Quantification Modesl

If the long read RNA-seq data consist of reads that always span the whole length of isoforms from which they are sequenced, then isoform expression quantification is a straightforward task, since where will be no multi-mapped reads. In such a case, the estimate for the relative abundance of an isoform is just the fraction of the sequenced reads that map to this isoform. However, although it is expected that the long reads have the same length as the isoforms sequenced (i.e. that the reads represent full-length transcripts), most of the these reads are, in fact, shorter than the isoforms from which they originate from due to technical artifacts, unexpected errors, fragmentation or breakage during sequencing, base-calling error, or for other reasons[4]. Additionally, the sequenced reads may constitute the full length of the sequenced molecule, but may not match the full length of the annotated isoform, because of well-known biological processes, such as transcript degredation.

The length distribution model for the long read RNA-seq dataset, sequenced from the Hct116 cell line (as discussed in Section 3.1), is presented along with the distribution of aligned lengths of the reads to the isoform and the distribution of isoform lengths in Figure S1. Here, we observe that the typical length of isoforms are 1.5 times longer than the typical length of the aligned segments of the reads that map to these isoforms. Therefore, we also observe multimapped reads at a non-trivial rate 70–80% in long read RNA-seq alignment, and probabilistic models and inference techniques like those applied to short read data [25] can also be useful for long read RNA-seq quantification. Although the EM algorithm in [25] can be used to quantify isoform expression from long read RNA-seq data, the generative model should, in fact, be modified, since long read RNA-seq experiments typically follow a protocol that is somewhat different than that followed in short read RNA-seq.

One such fundamental difference, on which we focus here, is that long read RNA-seq protocols do not include a systematic fragmentation step prior to sequencing. Specifically, since the fragment length of short read protocols is much less than the typical length of the the transcripts being sequenced, and since we wish to be able to generate reads from (nearly) any position within a transcript, these protocols almost universally contain a fragmentation step, in which the initial full-length transcripts are randomly fragmented, using one of several different techniques [20, 19].

The fundamental effect of this fragmentation process on the generative model is that we expect that the number of reads generated from a particular isoform to be proportional to the *product of* the number of copies of that isoform and the isoform’s length. For example, if two isoforms A and B were present in equal number in the underlying sample, but isoform A was *k* times longer than B, then we would expect, on average, to sequence approximately *k* times as many fragments from A as from B. This leads the number of reads generated from a particular isoform to have a fundamental dependence on the isoform’s length. However, fragmentation is typically not a step in long read RNA-seq protocols, so that we instead expect the number of reads arising from an isoform to be directly proportional to the number of copies of that isoform in the sample, and not, additionally, dependent on the isoform’s length. As we explain later, this seeming simplification to the model also eliminates a useful source of evidence when trying to determine the likely allocation of a multimapping read between sequence-similar transcripts of different length.

Several computational tools have been developed to quantify long-read RNA-seq data, but as discussed in Section 1, they typically remove the length effect without addressing the subsequent reduction it entails in the ability of the model to distinguish ambiguous fragments. Hence, we propose a model that removes the length effect of the short read model, and also replaces it with a new term that seeks to increase the uniformity of coverage under the read assignment probabilities.

### Proposed quantification framework

In this work, we refine the generative model introduced in [25] to enhance its applicability and accuracy specifically in the context of long read sequencing. The proposed generative model is illustrated in Figure 2 which can be useful to improve the likelihood function for long read RNA-seq quantification. The key insight behind our modification to the underlying model is that, in the short read sequencing model, the length dependence of the sequencing rate parameter provides useful information to help differentiate different potential origins of a read. For example, all other things being equal, a shorter transcript is a more likely origin for a read than a longer transcript, as the probability of selecting the specific fragment that starts at the position where the read aligns is higher for the shorter transcript than the longer transcript. Another way of viewing this is that the length parameter essentially penalizes uncovered stretches of transcripts where alignments do not occur. However, when we move to the long read model and remove this length dependence in the conditional probability of fragment generation, this discriminative component of the model is lost. This can, in turn, lead to counter-intuitive situations where the model fails to differentiate between potential mapping locations where a human observer may have a clear preference. See Figure 2 an illustrative example of such a case. At the same time, we do not want to include a length dependence in the long read quantification model, as we, in general, do not expect one given the underlying protocol and we expect that the length will not have a direct effect on the probability of sampling reads from a transcript — other than the effect that may arise from e.g. secondary and tertiary molecule structure and other biochemical preferences.

**Fig. 2.**
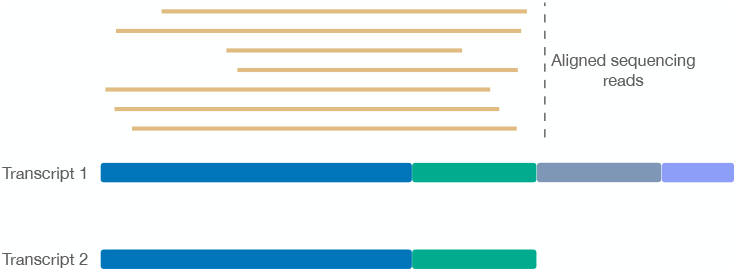
A toy example demonstrating how removing length effect from the probabilistic model can increase fragment assignment ambiguity in situations where human intuition may be clear. In this case, all of the sequenced reads align equally well to Transcript 1 and Transcript 2 (since Transcript 2 is, in fact, a proper prefix of Transcript 1). Given the totality of coverage, it is clear that these reads likely derive from Transcript 2, as the likelihood of generating this many reads from Transcript 1, but never sequencing beyond the second exon, is quite low. The model, however, does not encode this intuition, and does not convey a preference for Transcript 2 over Transcript 1 as an explanation for these reads.

Thus, we propose to incorporate a new term into the model which directly accounts for the potential coverage signal of the underlying transcripts. This term works to penalize large deviations in coverage over the body of a transcript when assessing the conditional probability of fragment assignment. This affords the model extra information to differentiate between potential allocations like those in Figure 2, and to prefer more intuitive explanations for the reads when they are available.

A plate diagram of our model is provided in Figure 1, where *R*_*n*_ and *CD*_*m*_ are the observed variables representing the long read sequences and potential coverage pattern for the *m*^*th*^ isoform, respectively. *θ* = [*θ*_1_,.., *θ*_*M*_] is the vector of isoform abundances that will be estimated via maximization of the associated likelihood. Additionally, *T*_*n*_, *A*_*n*_, and *SE*_*n*_ are the latent random variables representing the isoform, alignment score, and start-end positions of alignments respectively. The likelihood function for the modified generative model is given in Equation (2). Also, the explanation of the notations used in the generative model and likelihood function can be found inTable 1.

**Table 1.** Summary of Notation for Likelihood Function.

| Notation | Explanation |
| --- | --- |
| $\theta$ | Vector of isoform abundance parameters. |
| $M$ | Number of isoforms. |
| $N$ | Number of reads. |
| $T_n$ | Isoform random variable for read $n$. |
| $A_n$ | Alignment score random variable for read $n$. |
| $SE_n$ | Start-end position random variable for read $n$. |
| $CD_m$ | Coverage distribution model for isoform $m$. |
| $\mathcal{A}(r_n)$ | Set of all isoforms to which read $n$ aligns. |

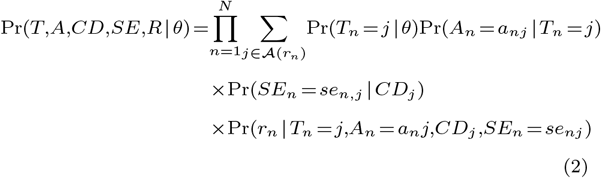

As seen in Figure 2, the (latent) random variable *T*_*n*_, denoting the transcript selected for read generation, only depends on the isoform expression parameters *θ* = [*θ*_1_,.., *θ*_*M*_]. In other words, if we know the expression level of all isoforms in the transcriptome, then the probability that we have selected the *j*^*th*^ isoform for sequencing is equal to its expression level or relative abundance *θ*_*j*_, so that Pr(*T*_*n*_ =*j*|*θ*)= *θ*_*j*_.

With the model in hand, the first step in the process of long read quantification is to align the reads to the target transcriptome to be quantified. For this task, we use `minimap2` [26, 27]. Of course, it is possible to make use of other aligners, or to make use of alignemnts of the read directly to the genome rather than to the transcriptome, but these are specific practical details that do not fundamentally affect the model being considered here. Throughout this work, we have adopted the `minimap2` parameters listed in section B of appendicies.

To obtain a reasonable model for Pr(*A*_*n*_ = *a*_*nj*_ | *T*_*n*_ = *j*), we use the alignment score computed by `minimap2` and encoded in the `AS` tag. Specifically, we consider all alignments of *n*^th^ read, and obtain the maximum alignment score among them. Let *j*^*0*^ be the alignment of the *n*^th^ read having maximum `AS`, which we denote as 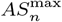. We then consider this conditional probability as 1, and model the probability of other alignments for this read to decrease exponentially as a function of the alignment score. Specifically, the function used to obtain Pr(*A*_*n*_ =*a*_*nj*_ |*T*_*n*_ =*j*) is given in Equation (3).

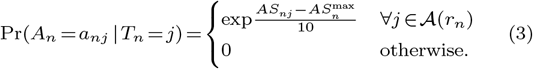

In Equation 3, *AS*_*nj*_ represent the alignment score of the *n*^*th*^ read aligned to *j*^*th*^ isoform, and *A*(*r*_*n*_) represents the set of all transcript indices to which read *r*_*n*_ aligns.

One pivotal insight derived from aligned reads to the transcriptome pertains to the observation and analysis of read coverage patterns specific to individual transcripts. As the distribution of transcript coverage pattern approaches uniformity, the likelihood of accurate alignments to that transcript appears to increase, consequently elevating the associated probability. The coverage pattern and uniformity varies substantially between different transcripts. For instance, Figure S2 shows the coverage patterns of three transcripts `ENST00000600659`, `ENST00000417088`, and `ENST00000449223` derived from aligning the Hct116 dataset (see Section 3) to the transcriptome. The coverage patterns in Figure S2 are extracted using IGV tools [43]. Specifically, SAMtools [12] was employed to acquire the depth of each position within the aforementioned three transcripts. The resulting data is depicted in Figure 3. It is evident, as observed in Figures S2 and 3, that `ENST00000417088` exhibits a coverage pattern closer to uniformity. In contrast, transcripts `ENST00000600659` and `ENST00000449223` display substantially less uniform patterns.

**Fig. 3.**
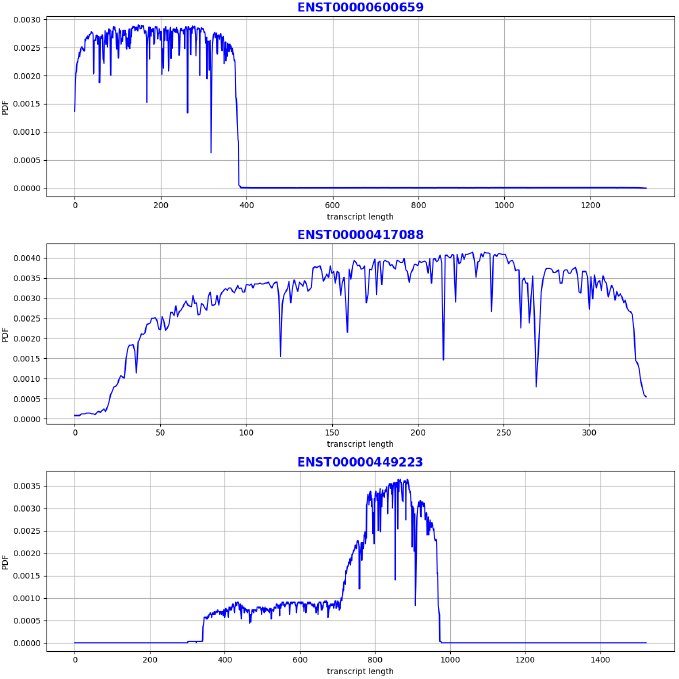
The coverage distribution model for three transcripts `ENST00000600659`, `ENST00000417088`, and `ENST00000449223`, as estimated from the BAM file of the Hct116 cell line dataset sequenced with direct cDNA.

Overall, our approach will be to attempt to incoporate the coverage distribution itself into the allocation probability for each fragment (that is, when attempting to determine the probability of each latent variable that assigns each alignment as the true origin of the read). This idea itself is quite general, and we believe that it can be explored and investigated thoroughly to determine the best way to make use of the coverage profiles in read allocation. However, here we propose one, specific, relatively simple model, and demonstrate that it helps to improve the quantification accuracy obtained by the model.

One approach to enhance the uniformity of these coverage patterns involves assigning to each alignment, a conditional probability that is related to the coverage pattern along the transcript being considered. That is, when the coverage pattern is non-uniform, and we are considering the alignment of a read to a highly-covered section of the transcript that would increase that non-uniformity, we would like to decrease the conditional probability of this alignment (assuming that there are other locations compatible with the read, where allocating this read would contribute less to non-uniformity of coverage). Likewise, when we are considering the alignment of a read to a sparsely-covered section of the transcript, we may wish to increase the conditional probability of alignment, to attempt to bring the coverage at the position covered by the read into closer concordance with the coverage of the rest of the transcript. The goal of such modifications to the conditional probabilities is to promote increased uniformity of coverage under the final allocations of reads. To accomplish this, first, we propose a model to compute a term inversely proportion to the coverage pattern for each transcript, which we elaborate below.

The process for acquiring the coverage distribution model for each transcript comprises the following six steps:

### Alignment

Align the reads to the transcriptome with `minimap2`. **Segmentation**: Segment the transcript into disjoint intervals. The number of segments is adjustable (default of 10 per transcript).

### Count reads

Count the number of reads that partially or completely cover each segment. The count for each read present in a segment is computed as the fraction of the segment length that is covered by that read. Specifically, the count for *i*^*th*^ segment is given by 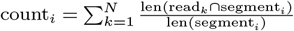, where *N*, read_*k*_, and segment_*i*_ represent the number of reads overlapping the *i*^*th*^ segment, the *k*^*th*^ read overlapping the *i*^*th*^ segment, and the *i*^*th*^ segment itself, respectively.

### Compute Probability

Compute the modified Binomial probability for each segment. Binomial probability is typically applicable to integer values, as the Binomial coefficient in its equation involves factorials, which inherently operate on integers. Here, we modify the Binomial probability to apply to real numbers. To achieve this modification, we use the Gamma function instead of the factorial function in the Binomial coefficient. The Gamma function serves as an extension of the factorial function, and can be evaluated on all complex numbers. The modified Binomial probability within the *i*^*th*^ segment is given in Equation (4), where *k*_*i*_ = count_*i*_, as computed above, and *n* is the summation of *k*_*i*_ over all segments of the transcript, given by 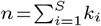, where *S* signifies the total number of segments.

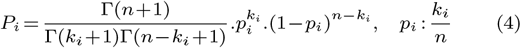

### Normalization

Normalize the modified Binomial probabilities for each segment on the transcript to guarantee that the sum across all segments in a transcript equals one.

### Discrete-Probability

Obtain the probability at each point on the transcript length. If we denote *P*_*i*_ as the computed probability for segment *i*, the probability assigned to each position within the segment *i* is determined by dividing the probability of the entire segment *i* by its corresponding length, as illustrated in Equation (5).

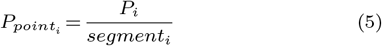

To explore what this obtained probability looks like across transcripts in informative examples, we plot the modified Binomial probability across the transcript length for three transcripts `ENST00000600659`, `ENST00000417088`, and `ENST00000449223` derived from Hct116 BAM file, as it is shown in Figure 4. As expected the probability evaluated across each of the mentioned transcripts is roughly inversely proportional to the coverage density shown in Figure S2 and Figure 3.

**Fig. 4.**
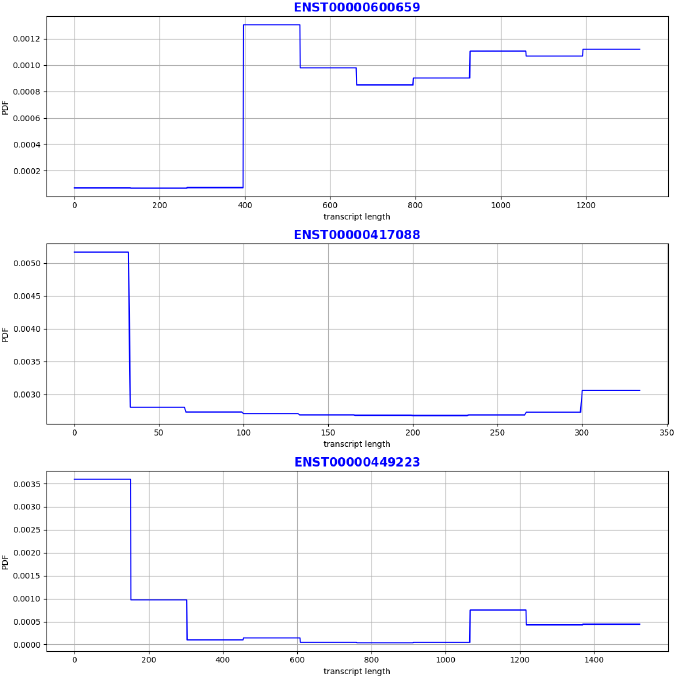
Modified binomial distribution for three transcripts `ENST00000600659`, `ENST00000417088`, and `ENST00000449223` from the Hct116 dataset BAM file.

Finally, with these modified coverage probabilities in hand, we can compute the probability Pr(*SE*_*n*_ = *se*_*nj*_ | *T*_*n*_ = *j*) the coverage probabilities computed across the corresponding transcript. This involves performing a summation over the modified Binomial probability across the positions ranging from the start to the end of the alignment on the transcript. The resulting probability is defined as 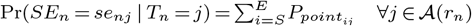. where *S* and *E* denote the start and end positions of the alignments across the transcript length, and 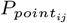 denotes the modified binomial probability at position *i* along the length of transcript *j*.

Given this definition of the relevant probabilities and conditional probabilities, we apply the Expectation-Maximization (EM) algorithm to derive the isoform expression level with the highest likelihood.

## Results

In this section, we conduct a detailed evaluation of our methodology using three distinct datasets, as listed in Section 3.1. Our analysis includes a comparison against alternative long read-centric quantification methodologies, namely `Bambu` [11], `LIQA` [21], `NanoCount` [17], and `ESPRESSO` [15]. To facilitate a rigorous comparison, we employ identical datasets and annotations as inputs for each methodology, and subsequently compute the Spearman correlation and Root Mean Square Error (RMSE) metrics for both the synthetic spike-in and sequin datasets and experimental datasets, obtained through ONT (and PacBio) sequencing technologies.

### Dataset

We employ three distinct datasets to assess and compare our model with other relevant methods. These datasets originate from three different cell lines including Hct116, H1975, and SH-SY5Y which are introduced in the Singapore Nanopore Expression Project [10], Long and short-read transcriptome profiling of human lung cancer cell lines [14], and the work of Wang et al.[45] introducing TEQUILA-seq, respectively. The reason behind selecting each of the mentioned dataset are discussed in section A of the appendices. Subsequently, all datasets are aligned to the genome and transcriptome as discussed in the Section B of the appendices.

### Linear Correlation Analysis

In order to assess the comparative performance of our proposed methodology against existing methods, we first evaluated the strength and direction of the linear relationship between isoform expression levels derived from long-read RNA-seq and short-read RNA-seq measurements of matched samples. This involved the generation of a scatter plot and determination of a best-fit regression line, providing a visual representation of the overall trend within the scatter plot. Additionally, the Pearson correlation coefficient is computed to quantitatively measure the strength of the linear relationship between isoform expression levels obtained from long-read RNA-seq and short-read RNA-seq measurements.

Figure 5, Figure S3 and Figure S4 display these results on the Hct116 cell line dataset [10] across the various quantification methods. Across all methods, the Pearson correlation p-values consistently approach zero, underscoring the statistical significance and robustness of the identified correlations. In Figure 5, the scatter plot captures the linear correlation pattern across all transcripts in the Hct116 cell line dataset. Our proposed method, with and without the coverage model (`oarfish_binomial` and `oarfish_NoCoverage`, respectively), along with comparative methods `Bambu` [11], `NanoCount` [17], and `ESPRESSO` [15], exhibit quite similar linear correlation patterns. Notably, `LIQA` [21] exhibits a markedly lower Pearson correlation (0.32), indicative of a weaker linear association. Moreover, our proposed method with coverage demonstrates an improvement (albeit slight) in Pearson correlation compared to the next best model, `ESPRESSO`.

**Fig. 5.**
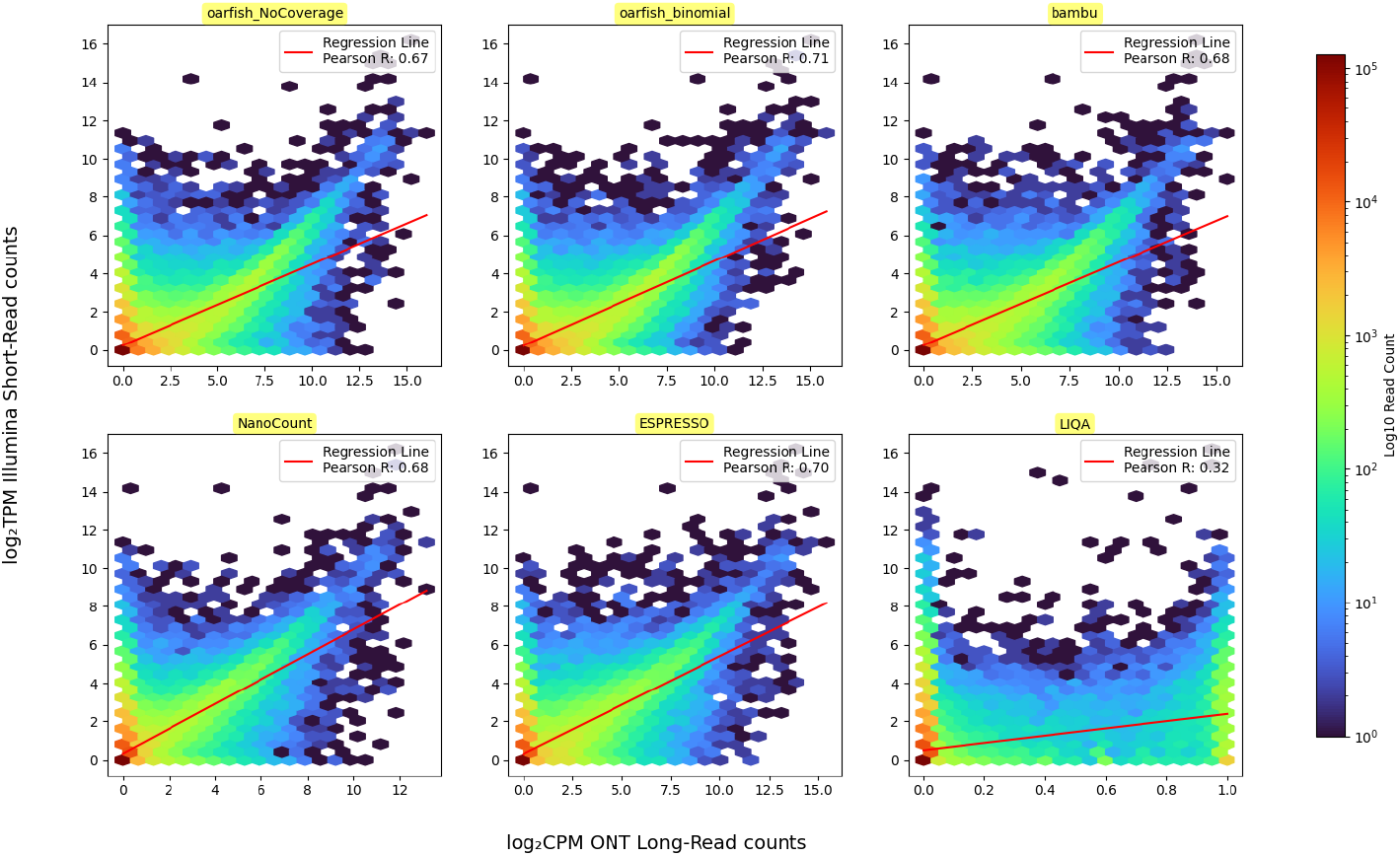
Scatter plot for Illumina short read counts and ONT long read counts for all the transcripts within the Hct116 cell line. The ONT long read RNA-seq dataset sequenced with direct cDNA protocol. The Illumina short read RNA-seq is quantified by Salmon. In all of these methods, the p-value for the Pearson correlation is almost zero (P-value *‘* 0.0). In this figure, the `oarfish_NoCoverage` and `oarfish_binomial` labels denote our proposed method without and with the binomial coverage model, respectively.

For further investigation, we generated two distinct scatter plots focusing on specific, previously identified subsets of important transcripts within the Hct116 cell line dataset, provided in [10]. Figure S3 provides an analogous scatter plot, but focusing exclusively on major transcripts. The observed pattern mirrors that of the plot featuring all transcripts in Figure 5, with a notable difference being the prevalence of a linear correlation among major transcripts, especially those with high abundance, across all methods except `LIQA`. Our proposed method with the binomial coverage model still exhibits a slight improvement (approximately 2%) compared to the next best model, which, in this case, is `NanoCount`. Additionally, the scatter plot focusing solely on sequin transcripts in the Hct116 cell line, as depicted in Figure S4, demonstrates a strong linear correlation between short and long read counts. This finding suggests that synthetic data, such as sequin transcripts, remains relatively unaffected by coverage models or other factors introduced in our proposed method or comparative models. It also suggests that such transcripts appear generally *easier* to accurately quantify compared to more complex and naturally occuring “biological” transcripts.

For extended analysis, we present scatter plots comparing short reads and long reads sequenced from H1975 [14] and SH-SY5Y [45] cell lines. Due to the substantial dataset sizes for H1975 (approximately 14 and 38 GB, respectively), the `ESPRESSO` and `LIQA` methods are excluded from the analysis due to their extended runtime, exceeding 48 hours. Figure S6 and Figure S8 illustrate that our proposed binomial coverage-based model again exhibits an improvement (approximately 3%) in Pearson correlation between short and long reads compared to the next best model. Notably, in the scatter plot for synthetic spike-in sequin transcripts in the H1975 cell line (Figure S6), a strong linear correlation between short and long read counts is observed. This finding aligns with our earlier observation of sequin transcripts abundance correlations in the Hct116 cell line.

### Non-linear Correlation and RMSE Analysis

In the previous section, we examined methods primarily based on the strength of linear correlation between short read and long read counts. To provide a more comprehensive evaluation, this section assesses the results in terms of the Spearman correlation and Root Mean Square Error (RMSE) as complementary metrics of accuracy. First, we ananlyze the result for the synthetic spike-in sequin transcripts with known concentrations in the Hct116 cell line. As depicted in Figure S5, all methods exhibit nearly identical Spearman correlation values for sequin long reads sequenced with the direct cDNA protocol, except for `LIQA`. However, under the RMSE, our proposed method with the binomial coverage model `oarfish_binomial` demonstrates a slightly smaller value than other methods, indicating a reduced RMSE (i.e. better performance). Conversely, the Spearman correlation for reads sequenced with directRNA is considerably lower compared to direct cDNA sequenced reads. The Spearman correlation for directRNA sequenced reads remains consistent across all methods, except for `Bambu` and `LIQA`, while the RMSE largely shows uniformity across all methods. Given that the synthetic spike-in sequin dataset has a known concentration, the results in Figure S5 underscore the commendable performance of our proposed method in terms of Spearman Correlation and RMSE. Furthermore, the slight changes in Spearman correlation and RMSE among different methods support the idea discussed in Section 3.2. These results further reiterates that coverage models, and other model adjustments like those adopted across other methods, seem to have little impact on performance on synthetic transcripts like sequins.

Next, we compared Spearman correlation and RMSE across methods for both all transcripts and for major transcripts in the Hct116 cell line. The Spearman correlation results indicate that our proposed method with the binomial coverage model `oarfish_binomial` performs well, along with `NanoCount`. Interestingly, NanoCount slightly outperforms others models for direct cDNA sequenced long reads. However, the scenario changes for major transcripts, where our proposed `oarfish_binomial` model shows a substantial 9% increase in Spearman correlation compared to the next best model. In terms of RMSE, the `oarfish_binomial` model consistently demonstrates superior performance, particularly when considering only major transcripts in the Hct116 cell line. Across all transcripts in the Hct116 cell line, our proposed model achieves the lowest RMSE for direct cDNA-sequenced long reads. For direct RNA-sequenced long reads, the RMSE remains similar across different methods. These results highlight the effectiveness of our proposed model, especially when focusing on the major transcripts in the cell line.

We performed the same analyses for the H1975 and SH-SY5Y cell lines. Notably, `ESPRESSO` and `LIQA` were excluded from the H1975 datasets due to their extended run times (exceeding 48 hours) in these samples. In Figure S9 and Figure S10, the ONT data produce consistently more concordant measurements with short read RNA-seq data than do PacBio reads, across all methods, for H1975 datasets. Our proposed model, `oarfish_binomial`, exhibited superior performance in Spearman correlation and RMSE for both the ONT and PacBio reads.

In Figure S11, our proposed method `oarfish_binomial` outperformed other methods in terms of Spearman correlation and RMSE, excluding LIQA [21] and ESPRESSO [15] due to their extended run times. Additionally, the SH-SY5Y dataset sequenced with 1DcDNA and directRNA protocols demonstrated higher accuracy than those sequenced with TEQUILA-seq. Moreover, an increase in sequencing time in TEQUILA-seq led to enhanced accuracy, as depicted in Figure S11, where the dataset with 8-hour sequencing outperformed the 4-hour dataset. Overall, the H1975 and SH-SY5Y datasets accord with the evaluations in the Hct116 dataset, and support the generally superior performance of our proposed method, `oarfish_binomial`, compared to the other methods in terms of Spearman correlation and RMSE.

Finally, we note that we performed an additional analysis using a variant of `oarfish` to try to ascertain the degree to which the long read data might reasonably concord with the short read estimates (i.e. what is a data-dependent upper limit on the similarity of predictions across these technologies?). These are the results in the figures annotated with the `+ shr` suffix. Specifically, we first estimated the transcript abundances from the short read data using `salmon`, and then we used the short read transcript quantification estimates to *initialize* the EM algorithm in `oarfish`. *Critically*, we allowed initializing transcripts estimated to be absent from the sample according to the short read data to 0 in the `oarfish` optimization, the result of which is forcing a 0 abundance for any transcript that was estimated to have 0 abundance according to the short read data. Interestingly, as shown in section (a) of Figure S10, this resulted in a quite substantial increase (around 20% to 30%) in Spearman correlation in certain datasets. However, such an increase in concordance is unlikely obtainable from long reads alone, at least in the current data, as we noticed a non-trivial number of transcripts that were estimated to have 0 abundance according to the short read data (and therefore were estimated to have 0 abundance under this initialization of `oarfish`), but which has *unique* read evidence in the long read data (and, of course, many zero to non-zero switches are observed in the other direction as well, though those remain represented even using this initialization).

### Runtime and Memory Usage

We conducted an analysis of the time and memory requirements of our proposed approach, and compared our tool with other state-of-the-art approaches, namely `Bambu`, `NanoCount`, `ESPRESSO`, and `LIQA`, using benchmarks across the Hct116, H1975, and SH-SY5Y datasets. The corresponding time and memory benchmarks are depicted in Figures 7, S12, and S13.

**Fig. 6.**
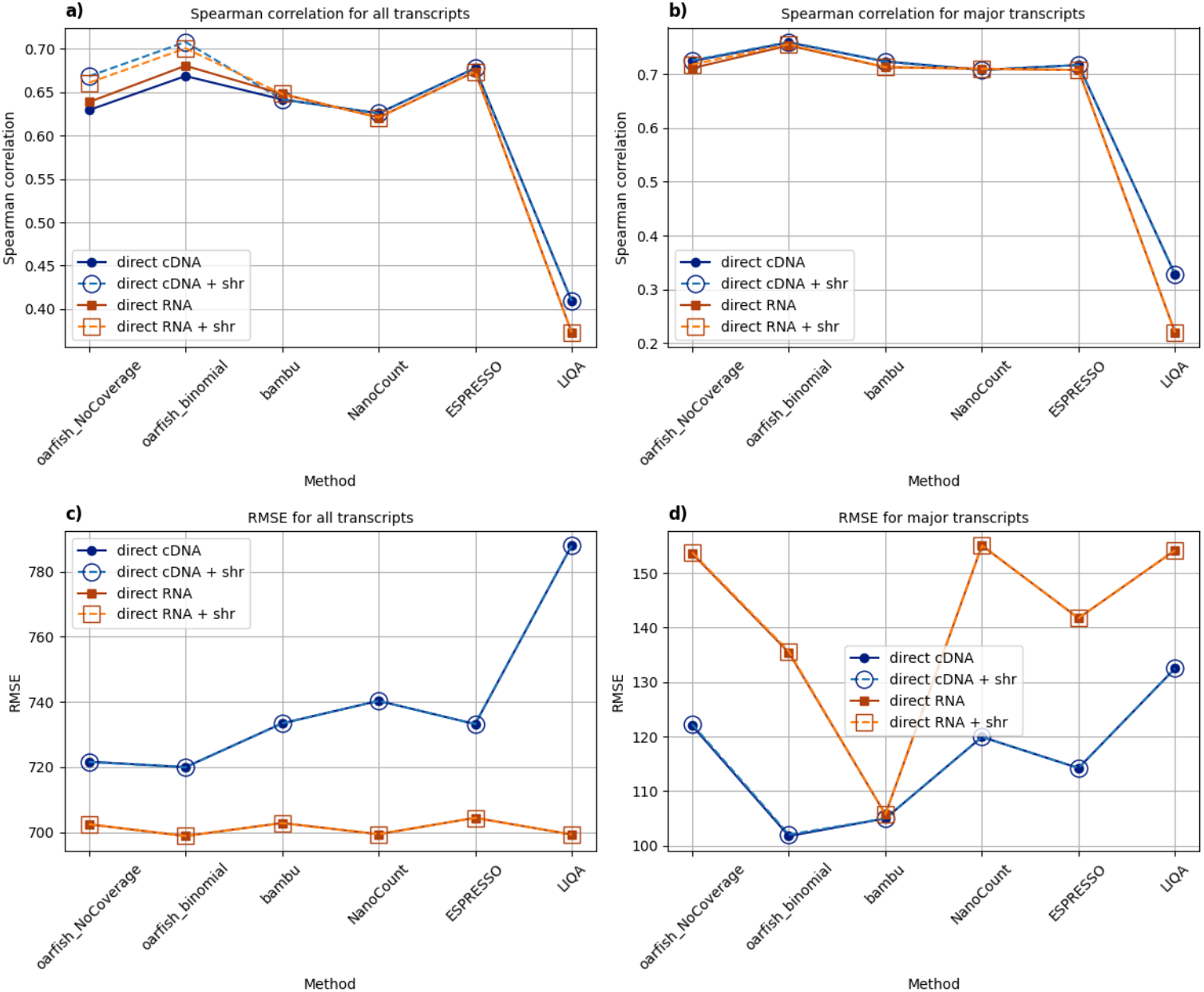
Spearman correlation and RMSE analysis for all transcripts and also major transcripts within in each gene in the Hct116 cell line: (a) Spearman correlation between Illumina short read and ONT long read RNA-seq datasets for all transcripts in Hct116 cell line. (b) Spearman correlation between Illumina short read and ONT long read RNA-seq datasets for only major transcripts in each gene within Hct116 cell line. (c) RMSE analysis for the all transcripts in the Hct116 cell line. (d) RMSE analysis for the only major transcripts in each gene within the Hct116 cell line. In these figures, `direct cDNA` and `direct RNA` represent ONT long reads sequenced with direct cDNA and direct RNA protocols, respectively. `shr` stands for short read counts, and `direct cDNA + shr` and `direct RNA + shr` indicate the use of Illumina short read counts for initialization in the `oarfish` EM algorithm during the quantification of transcripts in Hct116 cell line sequenced with direct cDNA and direct RNA protocols, respectively, in both proposed and comparative methods.

**Fig. 7.**
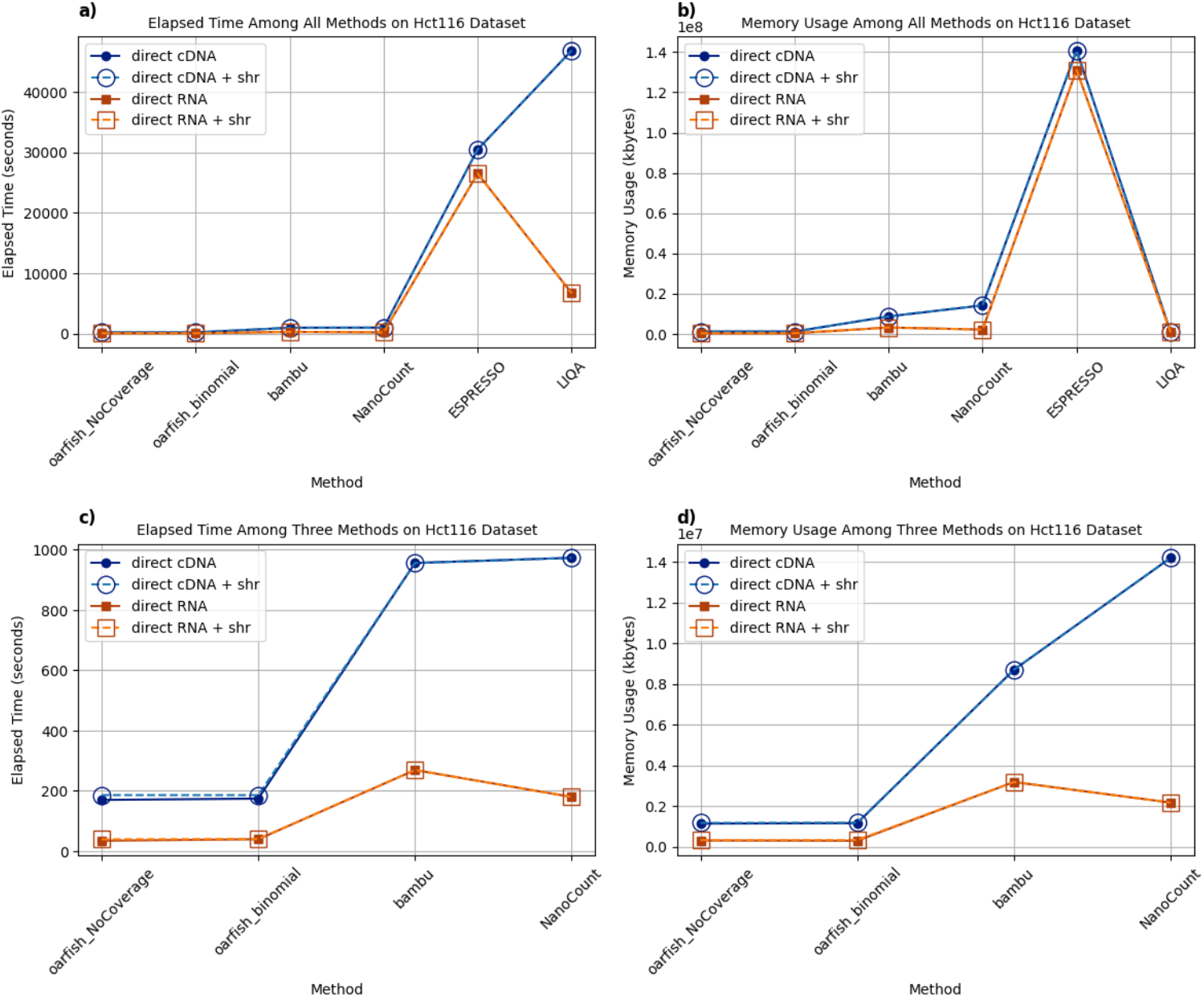
Performance metrics (elapsed time and peak memory usage) for the proposed and alternative methods using the Hct116 cell line datasets: (a) Elapsed time for all methods presented in seconds. (b) Memory usage for all methods displayed in kilobytes. (c) Elapsed time for `oarfish`, `bambu`, and `NanoCount` after excluding `ESPRESSO` and `LIQA`, providing a detailed comparison in seconds. (d) Memory usage for the same subset of methods, again excluding `ESPRESSO` and `LIQA`, enhancing the clarity of comparison and performance of specific tools, presented in kilobytes. In these figures, `direct cDNA` and `direct RNA` represent ONT long reads sequenced with direct cDNA and direct RNA protocols, respectively. `shr` stands for short read counts, and `direct cDNA + shr` and `direct RNA + shr` indicate the use of Illumina short read counts for initialization in the `oarfish` EM algorithm during the quantification of transcripts in Hct116 cell line sequenced with direct cDNA and direct RNA protocols, respectively, in both proposed and comparative methods.

Figure 7b and Figure S13b highlight notable variations in memory requirements among the tools. Specifically, `ESPRESSO` stands out with the highest memory demand (∼ 120GB to 140GB), while `oarfish` and `LIQA` exhibit the least consumption (only ∼ 1*GB*) for both Hct116 and H1975 datasets. Regarding elapsed time, significant distinctions emerge. While `oarfish`, `Bambu`, and `NanoCount` demonstrate efficient performance, `ESPRESSO` and `LIQA` run considerably slower, requiring several hours (Figure 7a and Figure S13a). As we have noted above, for the largest datasets, these methods did not complete within 48 hours, and were therefore excluded from evaluation.

To gain deeper insights, we excluded `ESPRESSO` and `LIQA` from the assessment, focusing on `oarfish`, `Bambu`, and `NanoCount`. These tools demonstrated substantially lower elapsed time and memory usage. Notably, `oarfish` showcased a marked reduction in both elapsed time and memory usage compared to the other two models across all datasets. This improvement becomes particularly pronounced as dataset sizes increase, as illustrated in Figure S12 for the H1975 dataset, as well as in the specific sequenced datasets, such as `direct cDNA` and `1D-cDNA` sequenced datasets from Hct116 and H1975 cell lines (Figure 7c, 7d, S13c, and S13d). In other words, `oarfish` improves elapsed time and memory usage about one order of magnitude for the largest sequenced dataset in each cell line. Moreover, as illustrated in Figures 7, S12, and S13, it is evident that employing short read counts in the EM algorithm exerts negligible impact on both execution time and memory usage, as is expected, since it only alters the initialization of the algorithm.

## Conclusion

In this paper we have introduced an enhanced model and tool for transcript quantification from long read RNA-seq data. Because of differences in the fundamental generative process (i.e. the absence of systematic fragmentation in long-read protocols and hence the lack of a length dependence in the abundance parameters), existing short-read quantification tools — at leas those that have not been explicitly retrofitted for long read quantification — are not well-suited to the task. At the same time, while numerous long read transcriptome analysis methods have been developed, most of these tools have prioritized identification over quantification. Yet, accurate quantification from long reads remains an unresolved challenge, and our emphasis here has been is on improving the quantification model itself, recognizing its significance alongside identification.

While existing long read quantification tools such as `Bambu`, `NanoCount`, `ESPRESSO`, and `LIQA` augment short-read quantification models with various additions, including read equivalence class categorization, distinct modifications of the read-isoform compatibility matrix, novel incorporation of read quality scores, and the read length distribution, none of these tools incorporate a model that explicitly accounts for the distribution of read coverage for individual transcripts. The read coverage pattern is a crucial factor, a piece of evidence that can shed light on the likelihood of sequenced reads originating from specific regions of transcripts. In our proposed model, we integrate a novel coverage distribution directly into the generative model that is used for transcript quantification. We have demonstrated that accounting for these coverage profiles can increase the accuracy of quantification from long-read RNA-seq data.

Additionally, an unsurprising but clear trend in the data we evaluated is that a higher read throughput can offer a more insightful perspective on the read coverage of transcripts, resulting in improved transcript expression estimation. This improvement is particularly highlighted in datasets obtained from high-throughput sequencing of the H1975 cell line, which was 20-40 times larger than datasets from Hct116 and SH-SY5Y cell lines. The outcomes demonstrate a significant enhancement in the quantification of the H1975 dataset. Notably, however, our proposed method, incorporating a coverage distribution model `oarfish_binomial`, exhibits performance improvements even in datasets with lower throughput, such as those sequenced from Hct116 and SH-SY5Y cell lines. This improvement is evident in terms of Pearson and Spearman correlation coefficients as well as RMSE, showcasing the robustness and versatility of our method across varying dataset sizes and sequencing depths. Furthermore, considering practical performance implications and computational resource usage, `oarfish` demonstrates highly-favorable behavior in terms of both execution speed and memory usage. This enhanced performance is particularly evident with larger dataset sizes, underscoring the scalability and efficiency of `oarfish`.

Although our proposed model improves quantification accuracy, and points at an interesting new direction for enhancing quantification models for long read data, it is but a first step and there are many potential directions for future work. Here, we highlight one theoretical and one practical direction for future work. From a theoretical perspective, our coverage model is one of the simpler models that one could imagine, based on a simple modified Binomial probability and, notably, *static* from the perspective of the inference algorithm. That is, we compute the coverage model given the initial alignments, and count a read as potentially covering *each* of its given multimapping locations. However, as we undertake inference, we learn meaningful information about where, among the many places from which a multimapping read might originate, the loci where it likely did originate. One interesting extension of our model would be to update the implied coverage profiles dynamically during inference based on the allocation probabilities for the reads, as they are inferred during the execution of the algorithm. This poses some of its own challenges (e.g. how to retain computational efficiency, and how to ensure convergence of the overall procedure). However, a similar approach is used to learn fragment length distributions [24] and bias models [37, 41] in existing short-read quantification tools, and a similar approach may be very promising in the context of the coverage distributions here.

As we have mentioned earlier, `oarfish` considers an alignment-level model for the purposes of optimization — that is, there is no summarization or aggregation of alignments into e.g. equivalence-classes [44, 38, 8, 37, 50]. As demonstrated by the time and memory efficiency of `oarfish`, this does not seem to pose a practical impediment to efficient implementation. Nonetheless, while existing standard factorizations of the likelihood would interfere with our coverage modeling, an interesting practical direction for future work is designing a factorization that is appropriate for application under our coverage model (or a *dynamic* variant of it). Such an approach could even further improve the computational efficiency of `oarfish`.

## Competing interests

R.P. is a co-founder of Ocean Genomics inc.

## Author contributions statement

Z.J. and R.P. devised the model and the experiments. R.P. and Z.J. both contributed to the `oarfish` implementation. Z.J. carried out the experiemnts and subsequent analysis. Z.J. and R.P. both wote the paper.

## Acknowledgments

This work has been supported by the US National Institutes of Health (R01 HG009937), and the US National Science Foundation (CCF-1750472, and CNS-1763680). Also, this project has been made possible in part by grant number 252586 from the Chan Zuckerberg Initiative Foundation. The founders had no role in the design of the method, data analysis, decision to publish, or preparation of the manuscript.

## Notes

https://github.com/COMBINE-lab/oarfish

