## Appendices for "Oarfish: Enhanced probabilistic modeling leads to improved accuracy in long read transcriptome quantification"

### Appendix A Dataset

The reason that we chose datasets sequenced from Hct116, H1975, and SH-SY5Y cell lines are as follows:

**The Singapore Nanopore Expression Project**[10]: We selected the dataset as it has been designed, collected, and distributed as a community resource meant to be ideal for aiding in method development and benchmarking. Additionally, the corresponding paper provides a useful classification concerning the major isoforms of each gene within each cell line, which can be used to "zoom in" on quantification estimates, and highlight differences in performance that may be otherwise hidden or obscured in transcriptome-wide assessments. This determination of major isoforms relies on the average expression levels for all isoforms associated with a gene, utilizing either long read or short read data. This specificity enables the assessment of most active isoforms individually, in addition to the evaluation of all isoforms collectively within each cell line. Furthermore, synthetic spike-in sequin data with specific concentrations has been incorporated into certain cell lines, including Hct116 (Colon cell line), K562 (Leukocytes cell line), and MCF7 (Breast cell line). This inclusion facilitates the assessment of isoform expression level quantification within an experimental dataset using external controls with known molar concentrations. In the subsequent analysis, we utilize both short read and long read RNA-seq datasets obtained from the Colon cell line, denoted as Hct116.

**Long and short-read transcriptome profiling of human lung cancer cell lines** [14]: Data from this study was selected due to the presence of a synthetic spike-in sequin control data, and also to the inclusion of sequenced long reads data using PacBio technology. The long read RNA-seq data obtained from PacBio should ideally exhibit higher base-calling precision and lower error rate compared to the dataset obtained from ONT technology [22], and should exhibit distinct artifacts or biases. Thus augmenting the dataset with long read RNA-seq data from PacBio sequencing, in conjunction with the short read RNA-seq data and long read RNA-seq from ONT, serves to establish a more robust ground truth for the comparative analysis of long read and short read data within each cell line. The datasets derived from pure RNA samples of Lung adenocarcinoma cell lines (H1975) are employed for additional assessment in this paper.

**TEQUILA-seq**[45]: TEQUILA-seq is a low-cost sequencing method implemented on the ONT platform which improves isoform coverage and quantification by mitigating the low throughput constraint of current long-read sequencing platforms. Consequently, datasets obtained from [45] serve as a valuable resource for demonstrating the applicability of our method, given its reliance on isoform coverage distribution. The improved precision in coverage distribution is important for enhancing the accuracy of results obtained from our proposed method. Additionally, the inclusion of synthetic spike-in SIRV data with known concentrations further augments the dataset's utility. This study exclusively utilizes samples derived from human neuroblastoma cell lines (SH-SY5Y), including both short read RNA-seq data and long read RNA-seq data obtained through three distinct sequencing methods: direct RNA, 1D cDNA, and TEQUILA-seq; all implemented on the ONT platform.

### Appendix B Alignment and Quantification Methods

#### B.1 Alignment Code

In this study, we utilized three distinct datasets derived from Hct116, H1975, and SH-SY5Y cell lines. The dataset originating from the Hct116 cell line was obtained through the Singapore Nanopore Expression Project (SG-NEx). Additionally, datasets from H1975 and SH-SY5Y cell lines were sourced from the Gene Expression Omnibus (GEO) under accession numbers GSE172421 and GSE213984, respectively.

To align the datasets to both the genome and transcriptome, we employed the following code. In the code, numbers 1 and 2 correspond to the alignment code used for ONT and PacBio datasets to the genome, while numbers 3 and 4 represent the code for aligning ONT and PacBio datasets to the transcriptome. The genome file hg38_sequins_SIRV_ERCCs_longSIRVs.fa

and transcriptome file hg38_sequins_SIRV_ERCCs_longSIRVs_cdna.fa used for this purpose were provided by the SG-NEx project. These files encompass synthetic SIRV and sequin transcripts, in addition to transcripts obtained from the GRCh38 reference genome. In the subsequent codes, variables such as `num_threads`, `genome_file`, `transcriptome_file`, `input_file`, and `output_file` represent the number of threads, genome file, transcriptome file, input fastq file, and output bam file, respectively:

1. `minimap2 -t num_threads -ax splice genome_file input_file > output_file`
2. `minimap2 -t num_threads -ax splice:hq -uf genome_file input_file > output_file`
3. `minimap2 -t num_threads -ax map-ont -N 100 transcriptome_file input_file > output_file`
4. `minimap2 -t num_threads -ax map-pb -N 100 transcriptome_file input_file > output_file`

#### B.2 Quantification Code

We employed `salmon` for quantifying short-read RNA-seq data. The procedure involved generating an index file and subsequently utilizing this index to quantify the provided fastq files. The code snippet used for this process is outlined below.

1. `salmon index -p num_threads -t transcriptome_file -i output_index -k 31`
2. `salmon quant -p num_threads -i output_index -l A -1 input_file_1 input_file_2 -o output_file`

We employed two distinct codes to quantify long reads with **oarfish**, one incorporating a coverage distribution model and the other without. These codes are outlined below.

1. `oarfish --alignments input_bam_file --threads num_threads --output-path output_directory --filter-group no-filters`
2. `oarfish --alignments input_bam_file --threads num_threads --output-path output_directory --model-coverage --filter-group no-filters`

We utilized the following code for quantifying the dataset using **bambu**.

```
1. bambuAnnotations = prepareAnnotations(gtf_annotation)
2. RcOut1 = bambu(reads = input_bam_file, annotations = bambuAnnotations, genome = genome_file, discovery = FALSE)
```

The subsequent code is utilized for quantifying datasets sequenced with direct cDNA and direct RNA, employing **NanoCount** for each respective dataset.

```
1. NanoCount -i input_bam_file --keep_neg_strand -d -1 -b output_bam_selected -o output_file
2. NanoCount -i input_bam_file -b output_bam_selected -o output_file
```

The subsequent code is employed for quantifying the dataset using **ESPRESSO**.

```
1. perl ESPRESSO_S -A annotation_file -F genome_file -L tsv_file -O output_directory -T num_threads
2. perl ESPRESSO_C -I output_directory -F genome_file -X 0 -T num_threads
3. perl ESPRESSO_Q -A annotation_file -L tsv_file -V output_file -T num_threads
```

To quantify the dataset with **LIQA**, we employed the following code.

```
1. liqa -task refgene -ref annotation_file} -format gtf -out -O output_refgene
2. liqa -task quantify -refgene output_refgene -bam sorted_input_bam_file -out output_file -max_distance 10 -f_weight 1
```

### Appendix C Supplementary Figures

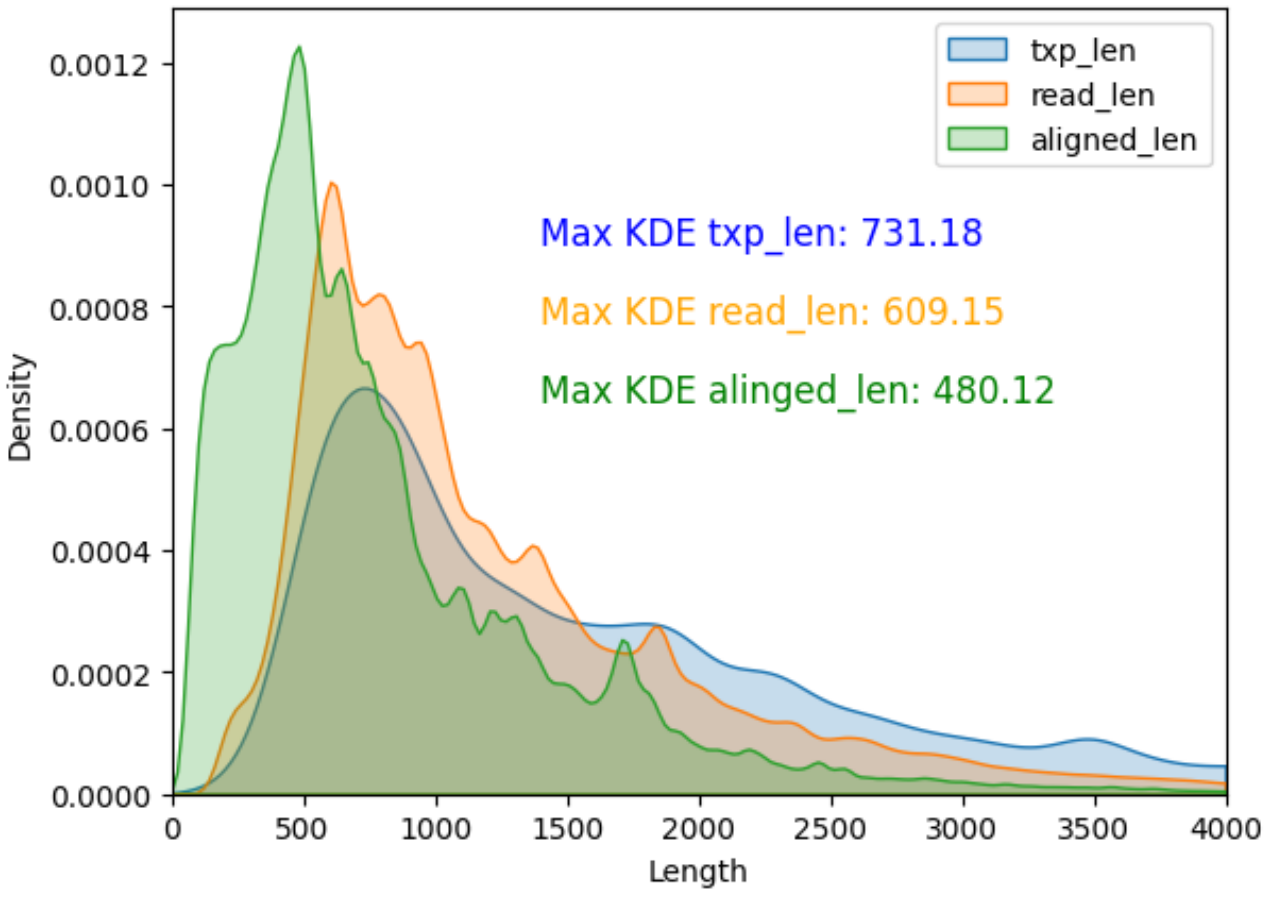


**Supplementary Figure S1.** Transcript length, read length, and aligned part of the read length density distribution. This figure illustrates the density distribution of transcript length (`txp_len`), read length (`read_len`), and aligned length of reads to transcripts (`aligned_len`). The analysis is conducted using a long-read RNA-seq dataset from the Hct116 cell line, sequenced with the direct cDNA protocol. The length distribution is depicted up to a maximum length of 4000.

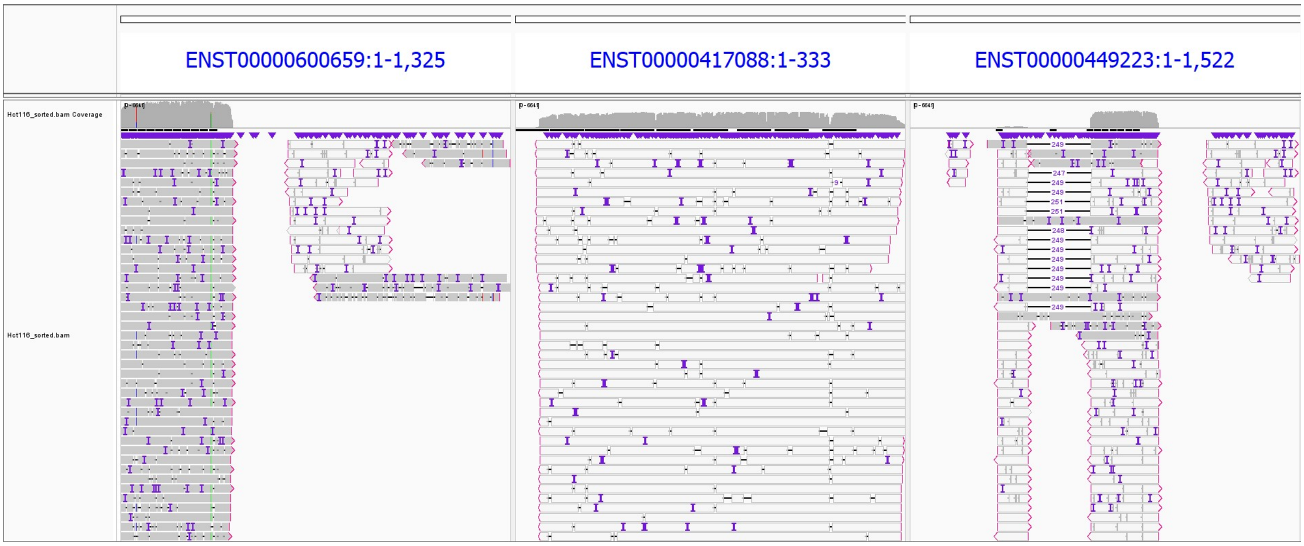


**Supplementary Figure S2.** The output of IGV tools for three transcripts `ENST00000600659`, `ENST00000417088`, and `ENST00000449223` from the BAM file of the Hct116 cell line dataset sequenced with direct cDNA.

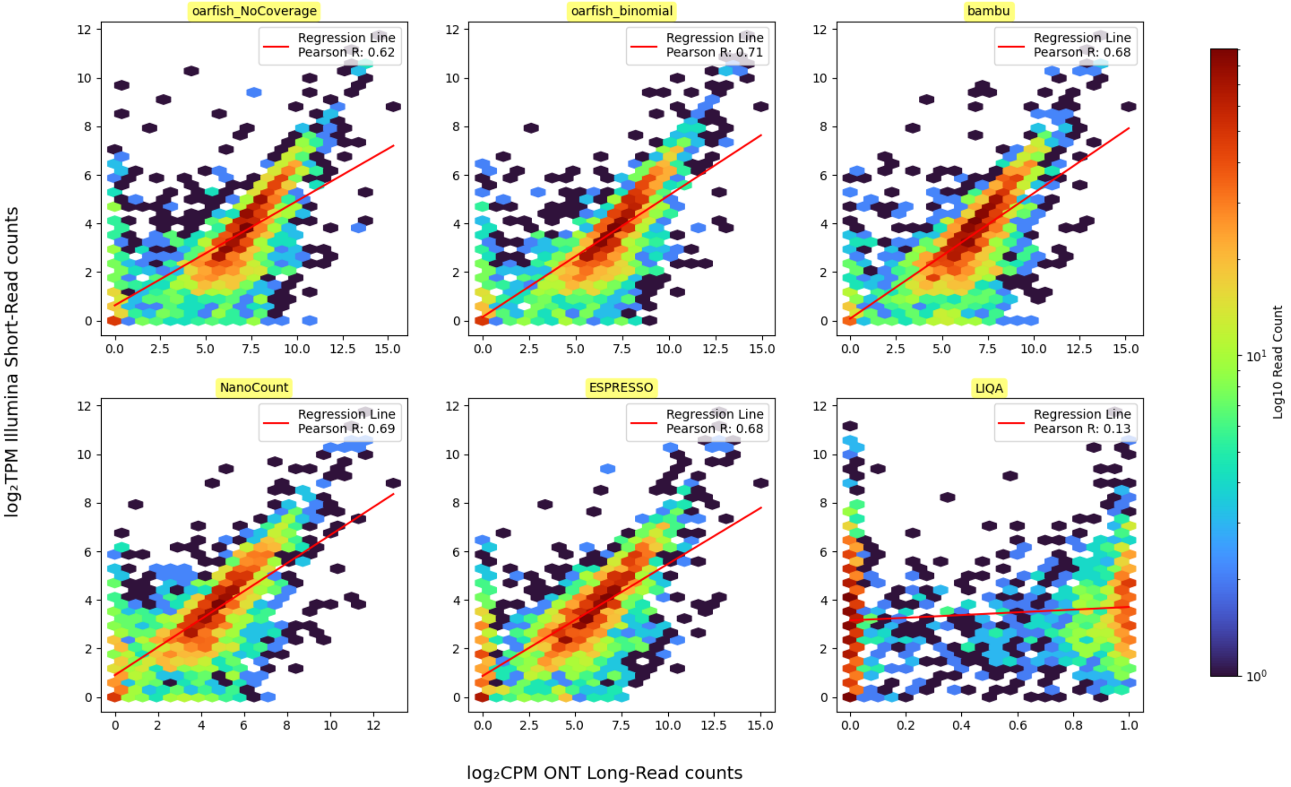


**Supplementary Figure S3.** Scatter plot for Illumina short read counts and ONT long read counts for only major transcripts in each gene within the Hct116 cell line. The ONT long read RNA-seq dataset sequenced with direct cDNA protocol. The Illumina short read RNA-seq is quantified by Salmon. In all of these methods, the p-value for the Pearson correlation is almost zero (P-value $\simeq$0.0). In this figure, the `oarfish_NoCoverage` and `oarfish_binomial` labels denote our proposed method without and with the Binomial coverage model, respectively.

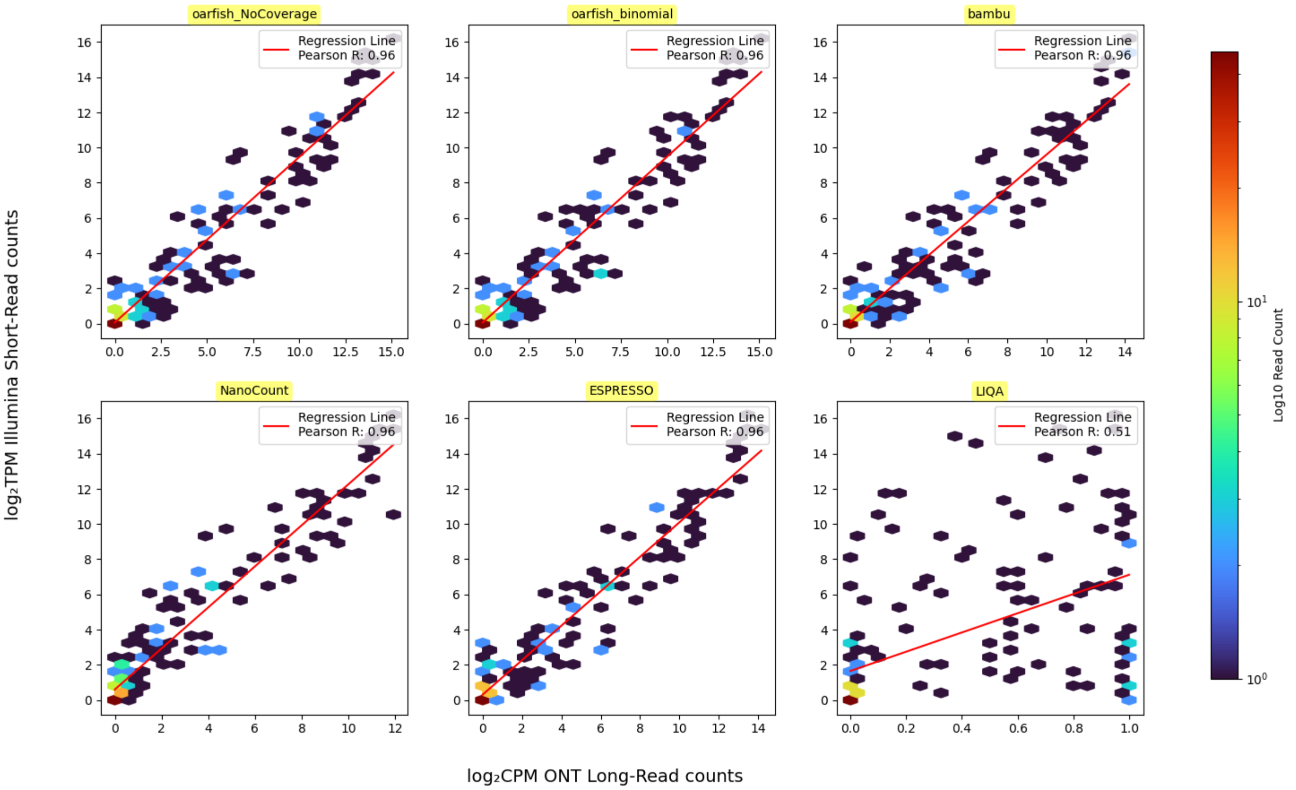


**Supplementary Figure S4.** Scatter plot for Illumina short read counts and ONT long read counts for only synthetic spike-in sequin transcripts in the Hct116 cell line. The ONT long read RNA-seq dataset sequenced with direct cDNA protocol. The Illumina short read RNA-seq is quantified by Salmon. In all of these methods, the p-value for the Pearson correlation is almost zero (P-value $\simeq$0.0). In this figure, the `oarfish_NoCoverage` and `oarfish_binomial` labels denote our proposed method without and with the binomial coverage model, respectively.

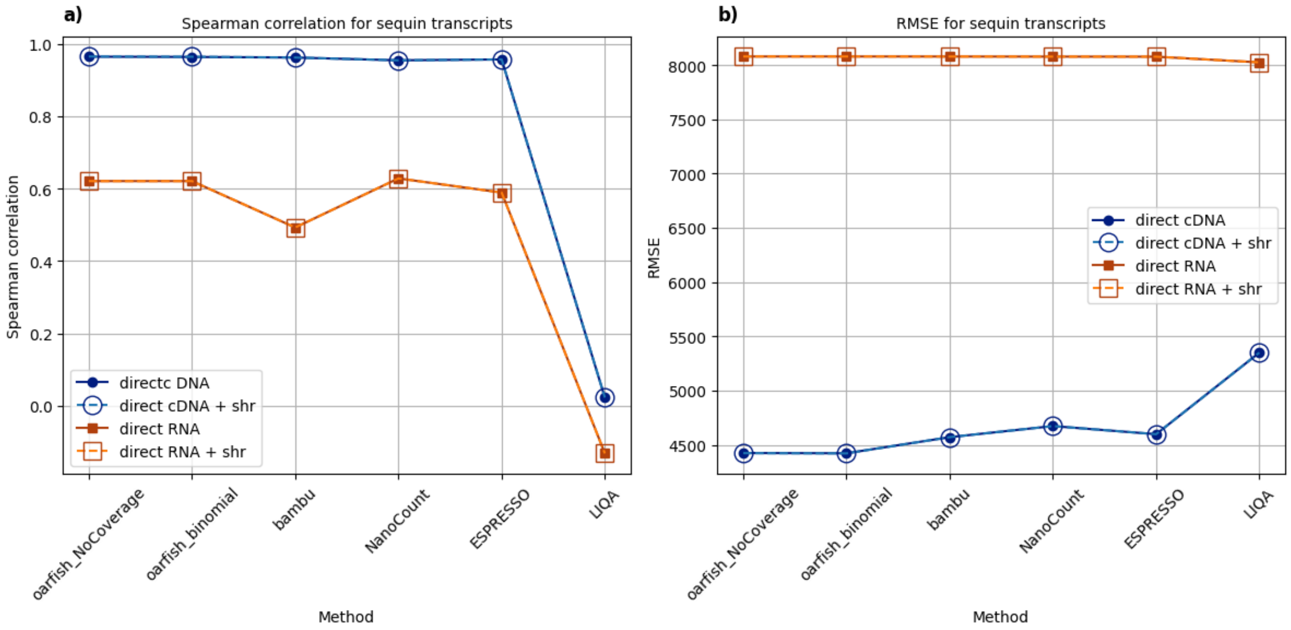


**Supplementary Figure S5.** Spearman Correlation and RMSE analysis for synthetic spike-in sequin transcripts in the Hct116 cell line: (a) Spearman correlation between Illumina short read and ONT long read RNA-seq datasets for synthetic spike-in sequin transcripts. (b) RMSE analysis for the same synthetic spike-in sequin transcripts. In these figures, 'direct cDNA' and 'direct RNA' represent ONT long reads sequenced with direct cDNA and direct RNA protocols, respectively. 'shr' stands for short read counts, and 'direct cDNA + shr' and 'direct RNA + shr' indicate the use of Illumina short read counts for initialization in the oarfish EM algorithm during the quantification of transcripts in Hct116 cell line sequenced with direct cDNA and direct RNA protocols, respectively, in both proposed and comparative methods.

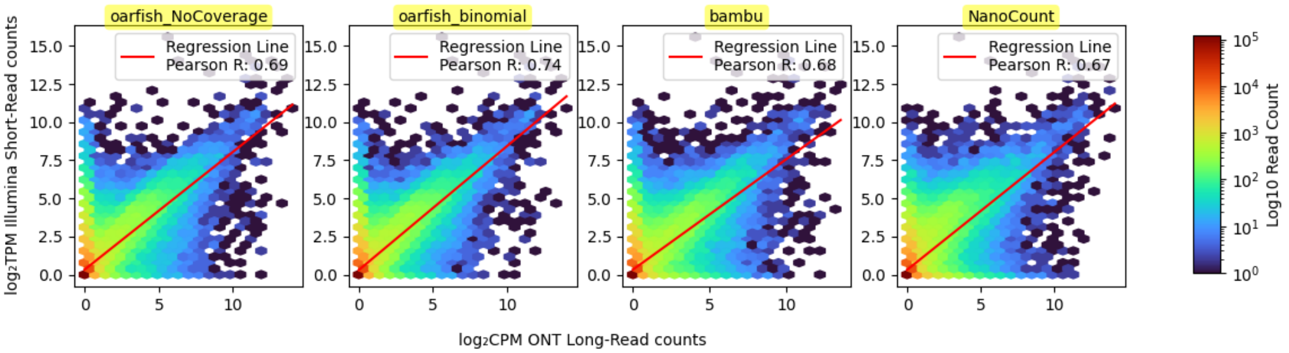


**Supplementary Figure S6.** Scatter plot for Illumina short read counts and ONT long read counts for all the transcripts within the H1975 cell line. The ONT long read RNA-seq dataset sequenced with direct RNA protocol. The Illumina short read RNA-seq is quantified by Salmon. In all of these methods, the p-value for the Pearson correlation is almost zero (P-value $\simeq$0.0). In this figure, the `oarfish_NoCoverage` and `oarfish_binomial` labels denote our proposed method without and with the binomial coverage model, respectively.

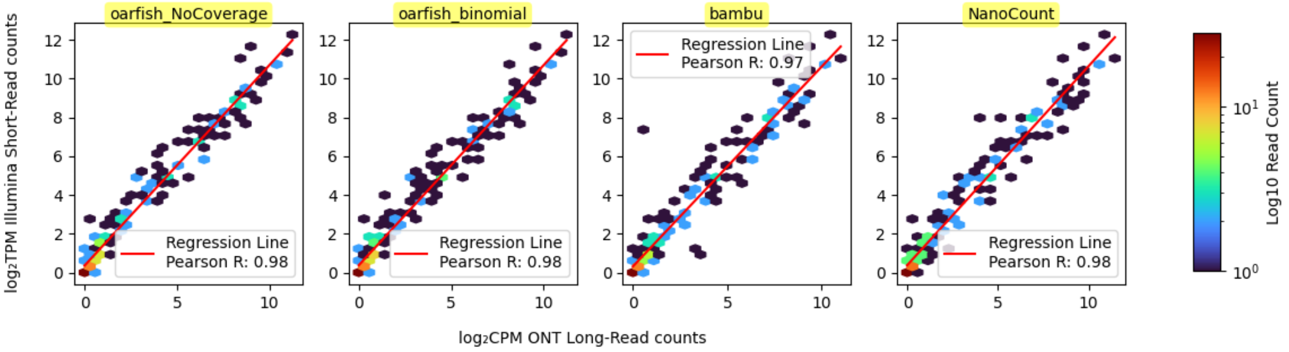


**Supplementary Figure S7.** Scatter plot for Illumina short read counts and ONT long read counts for only synthetic spike-in sequin transcripts in the H1975 cell line. The ONT long read RNA-seq dataset sequenced with direct RNA protocol. The Illumina short read RNA-seq is quantified by Salmon. In all of these methods, the p-value for the Pearson correlation is almost zero (P-value $\simeq$0.0). In this figure, the `oarfish_NoCoverage` and `oarfish_binomial` labels denote our proposed method without and with the binomial coverage model, respectively.

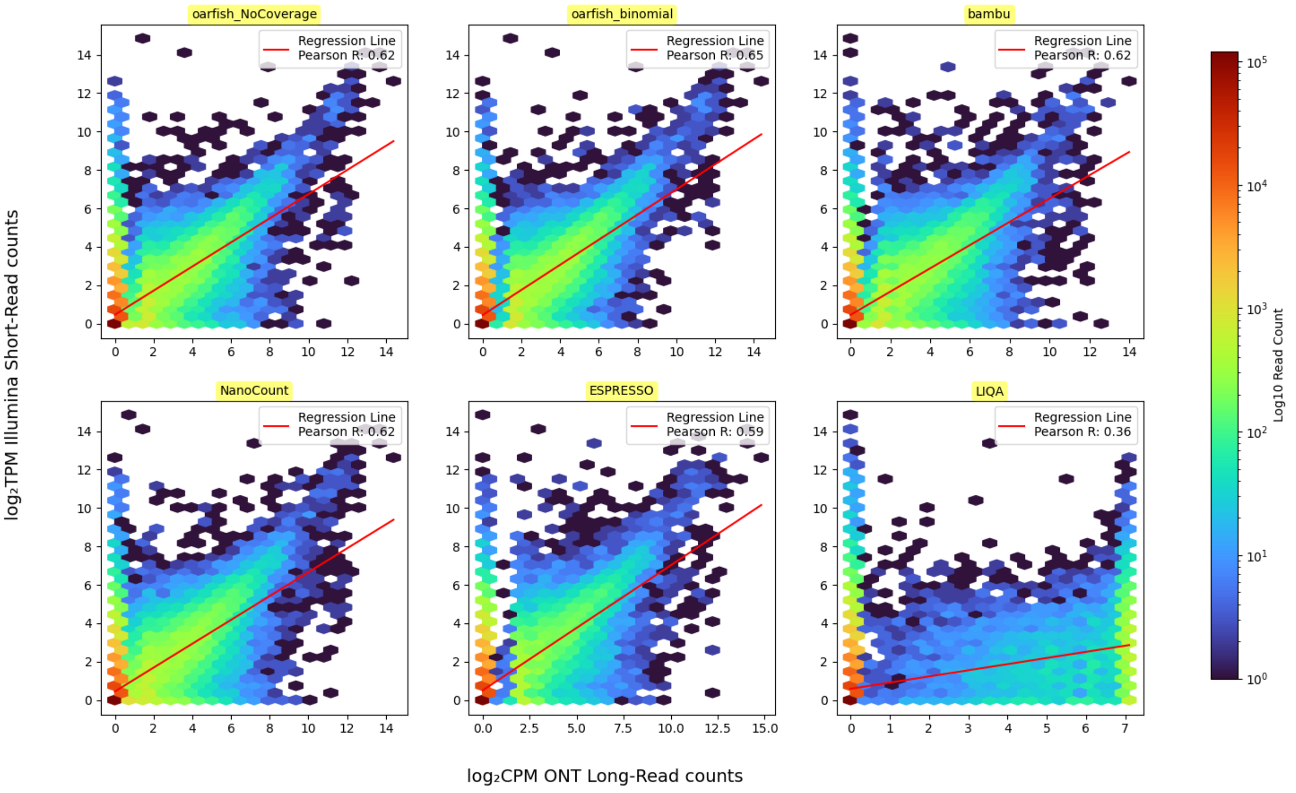


**Supplementary Figure S8.** Scatter plot for illumina short read counts and ONT long read counts for all the transcripts within the SH-SY5Y cell line. The ONT long read RNA-seq dataset used here is sequenced with direct RNA protocol. The Illumina short read RNA-seq is quantified by `salmon`. In all of these methods, the p-value for the Pearson correlation is almost zero (P-value $\simeq 0.0$). In this figure, the `oarfish_NoCoverage` and `oarfish_binomial` labels denote our proposed method without and with the binomial coverage model, respectively.

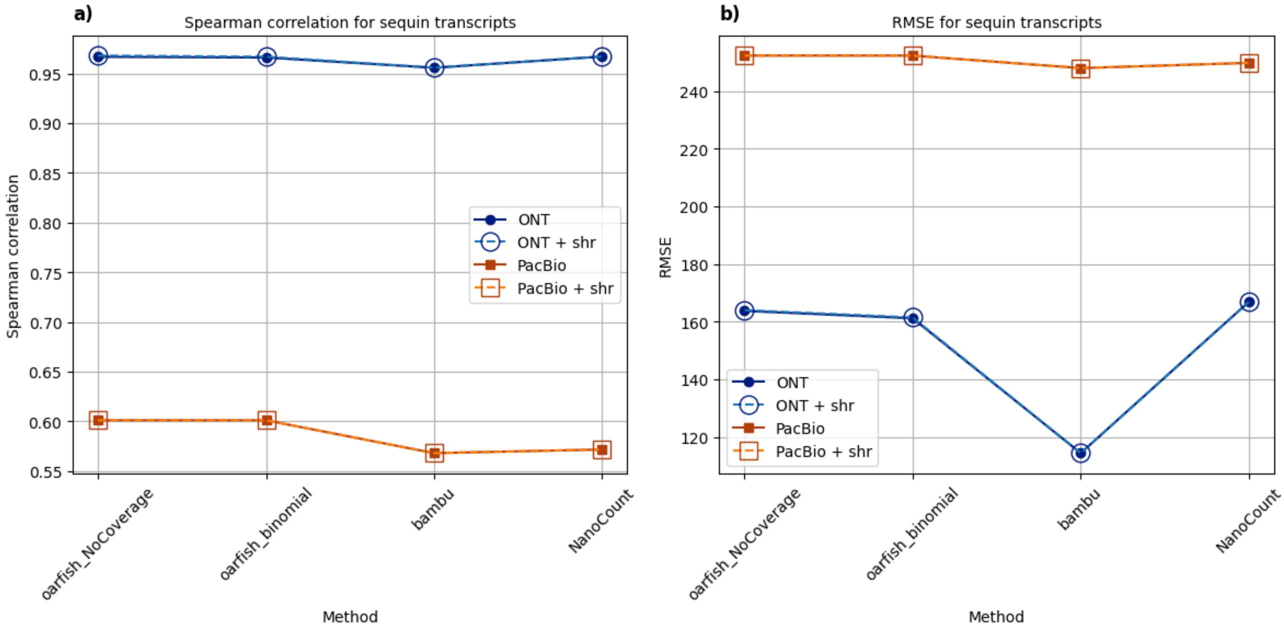


**Supplementary Figure S9.** Spearman correlation and RMSE analysis for synthetic spike-in sequin transcripts in the H1975 cell line: (a) Spearman correlation between Illumina short read and long read datasets obtained with ONT and PacBio for synthetic spike-in sequin transcripts. (b) RMSE analysis for the same synthetic spike-in sequin transcripts. 'shr' stands for short read counts, and `ONT + shr` and `PacBio + shr` indicate the use of Illumina short read counts for initialization in the `oarfish` EM algorithm during the quantification of transcripts in H1975 cell line sequenced with ONT and PacBio, respectively, in both proposed and comparative methods.

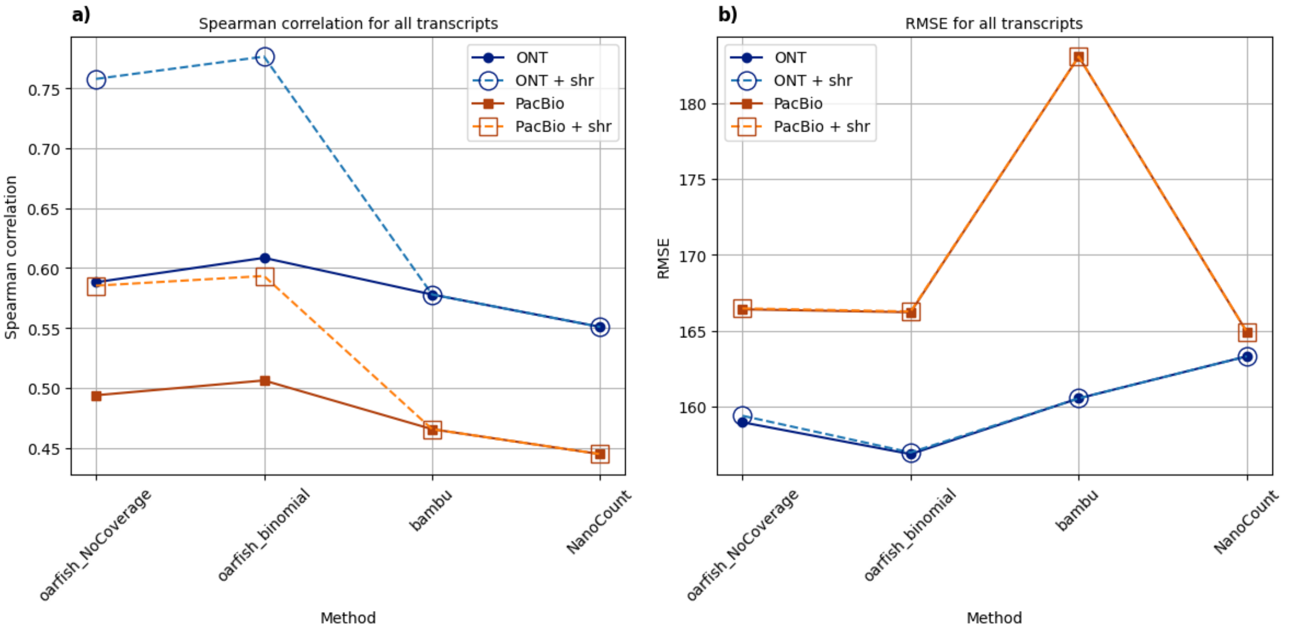


**Supplementary Figure S10.** Spearman correlation and RMSE analysis for all transcripts and also major transcripts in aach gene within the H1975 cell line: (a) Spearman correlation between Illumina short read and long read datasets obtained with ONT and PacBio for all transcripts in H1975 cell line. (b) RMSE analysis for the all transcripts in the H1975 cell line. `shr` stands for short read counts, and `ONT + shr` and `PacBio + shr` indicate the use of Illumina short read counts for initialization in the `oarfish` EM algorithm during the quantification of transcripts in H1975 cell line sequenced with ONT and PacBio, respectively, in both proposed and comparative methods.

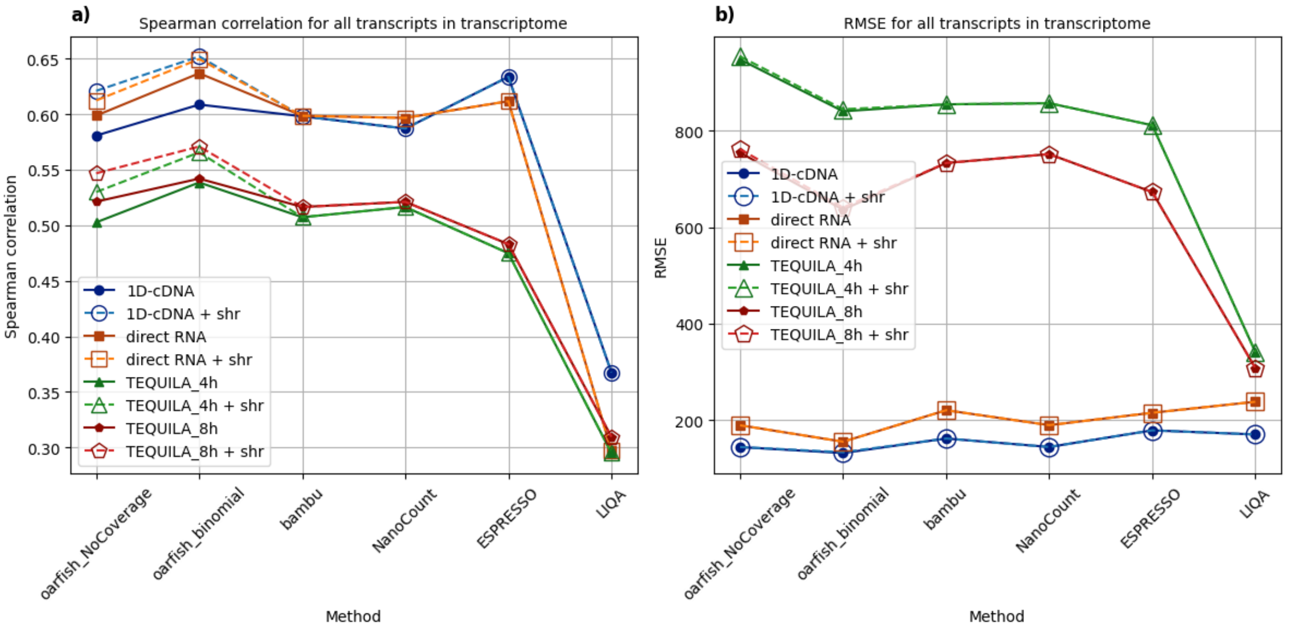


**Supplementary Figure S11.** Spearman correlation and RMSE analysis for all transcripts and also major transcripts in each gene within the SH-SY5Y cell line: (a) Spearman correlation between Illumina short read and ONT long read RNA-seq datasets for all transcripts in the SH-SY5Y cell line. (b) RMSE analysis for the all transcripts in the SH-SY5Y cell line. RMSE analysis for the only major transcripts in each gene within the SH-SY5Y cell line. In these figures, `1D-cDNA` and `direct RNA` represent ONT long reads sequenced with 1D-cDNA and direct RNA protocols, respectively. `shr` stands for short read counts, and `1D-cDNA + shr`, `direct RNA + shr`, `TEQUILA_4h + shr`, and `TEQUILA_8h + shr` indicate the use of Illumina short read counts for *initialization* in the `oarfish` EM algorithm during the quantification of transcripts in SH-SY5Y cell line sequenced with 1D-cDNA, direct RNA, and TEQUILA-seq protocols, respectively, in both proposed and comparative methods. `TEQUILA_4h` and `TEQUILA_8h` denote the TEQUILA-seq dataset with 4 hour and 8 hour sequencing time.

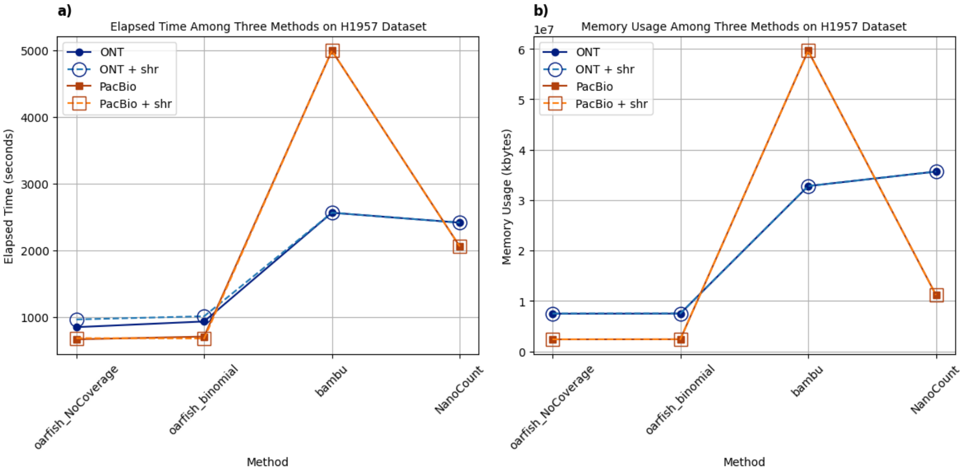


**Supplementary Figure S12.** Performance metrics for proposed and comparative methods using the H1975 cell line datasets: (a) Elapsed time for all methods presented in seconds. (b) Memory usage for all methods displayed in kilobytes. In these figures, `shr` stands for short read counts, and `ONT + shr` and `PacBio + shr` indicate the use of Illumina short read counts for initialization in the `oarfish` EM algorithm during the quantification of transcripts in H1975 cell line sequenced with ONT and PacBio, respectively, in both proposed and comparative methods.

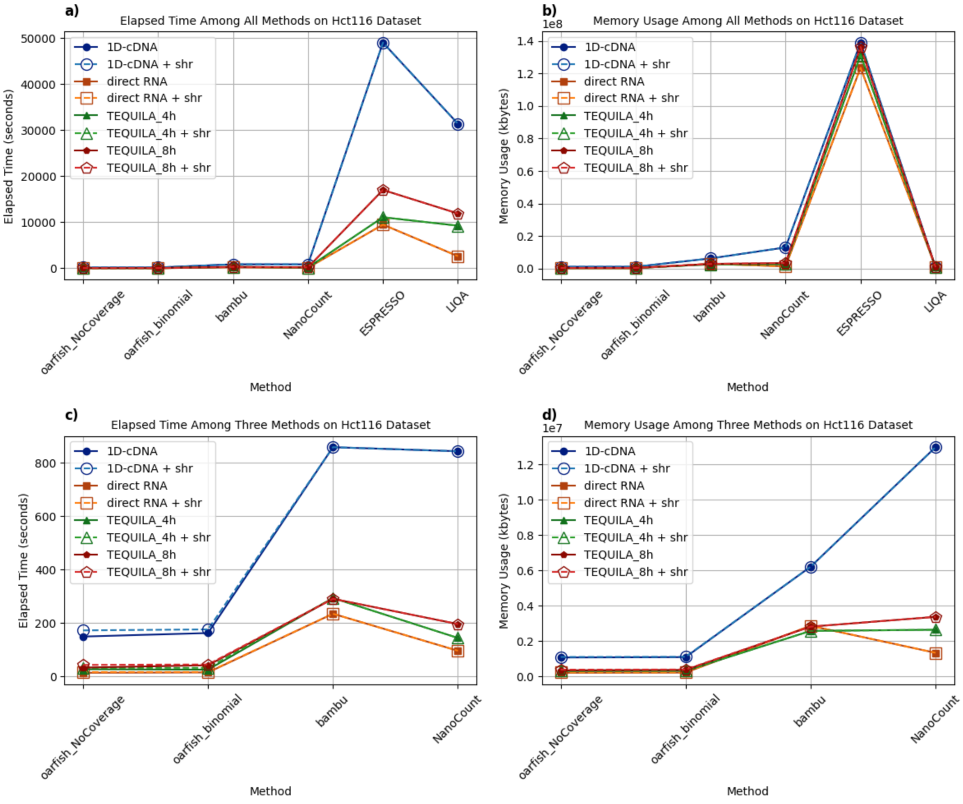


**Supplementary Figure S13.** Performance metrics for proposed and comparative methods using the SY5Y cell line datasets: (a) Elapsed time for all methods presented in seconds. (b) Memory usage for all methods displayed in kilobytes. (c) Elapsed time for `oarfish`, `bambu`, and `NanoCount` after excluding `ESPRESSO` and `LIQA`, providing a detailed comparison in seconds. (d) Memory usage for the same subset of methods, again excluding `ESPRESSO` and `LIQA`, enhancing the clarity of comparison and performance of specific tools , presented in kilobytes. In these figures, `1D-cDNA` and `direct RNA` represent ONT long reads sequenced with 1D-cDNA and direct RNA protocols, respectively. `shr` stands for short read counts, and `1D-cDNA + shr`, `direct RNA + shr`, `TEQUILA_4h + shr`, and `TEQUILA_8h + shr` indicate the use of Illumina short read counts for *initialization* in the `oarfish` EM algorithm during the quantification of transcripts in SH-SY5Y cell line sequenced with 1D-cDNA, direct RNA, and TEQUILA-seq protocols, respectively, in both proposed and comparative methods. `TEQUILA_4h` and `TEQUILA_8h` denote the TEQUILA-seq dataset with 4 hour and 8 hour sequencing time.
